# Deleterious effects of thermal and water stresses on life history and physiology: a case study on woodlouse

**DOI:** 10.1101/2022.09.26.509512

**Authors:** Charlotte Depeux, Angèle Branger, Théo Moulignier, Jérôme Moreau, Jean-François Lemaître, François-Xavier Dechaume-Moncharmont, Tiffany Laverre, Hélène Pauhlac, Jean-Michel Gaillard, Sophie Beltran-Bech

## Abstract

We tested independently the influences of increasing temperature and decreasing moisture on life history and physiological traits in the arthropod *Armadillidium vulgare*. Both increasing temperature and decreasing moisture led reproductive success to decrease. While the density of immune cells decreased and the β-galactosidase activity increased with increasing temperature and decreasing moisture, which suggests a negative impact of these stressors on individual performance, increased temperature and decreased moisture affected differently the other biomarkers conjuring different underlying mechanisms depending on the stress applied. Our findings demonstrate overall a negative impact of high temperature and low moisture on woodlouse welfare. Changing temperature or moisture had slightly different effects, illustrating the need to test further the respective role of each of these key components of climate change on organisms to predict more reliably the future of our ecosystems.

## Introduction

The Intergovernmental Panel on Climate Change (IPCC) forecasts an average increase in temperature between +1.5°C and +4°C in 2100 (Masson-Delmotte et al., 2021). Not only will average temperatures and the frequency and intensity of precipitation change, but extreme events will increase in frequency. Although the link between global warming and drought is still highly debated and may not be direct (Trenberth et al., 2014), droughts due to a decrease in rainfall and an increase in evaporation are expected to take place in the coming decades (Dai, 2013), and should be much more intense than current droughts (Trenberth et al., 2014). As deterioration of environmental conditions are known to impact life history traits such as growth rate, reproductive success, or longevity (e.g. Chen et al., 2019; Johnson and Jones, 2016; Khadioli et al., 2014), identifying the potential implications of climate change for organisms is a research question of paramount importance.

Terrestrial arthropods are ectotherms that are particularly sensitive to temperature and moisture changes (Lister and Garcia, 2018; Maron et al., 2015). Global warming constitutes a threat for them (Johnson and Jones, 2016). In Lepidoptera, for example, too high temperatures prevent hatching (Khadioli et al., 2014). In both Lepidoptera and Hymenoptera, increasing temperature beyond the optimum can have detrimental effect on survival (Abou-Shaara et al., 2012; Khadioli et al., 2014). In some Coleoptera, egg viability decreases and hatching time increases for viable eggs when they are exposed to drought (Johnson et al., 2010). When facing the costs of increased temperature and drought frequency on life history traits, arthropods display different responses to resist or tolerate such changes (Strachan et al., 2015). For example, some arthropods can migrate to refugia, others can implement physiological resistance tactics (e.g. resistant eggs) and/or dormancy, and others are able to alter their life cycle and/or development (Strachan et al., 2015; Verberk et al., 2008). In organisms with limited movement capacity, increased temperature and decreased moisture are expected to induce pronounced physiological stresses. Studying how these stresses affect both life history and physiological traits would allow us to anticipate the effect of global warming on organisms with limited movement capacity.

The common woodlouse *Armadillidium vulgare* is a key soil decomposer naturally exposed to a wide range of environmental conditions (Souty-Grosset et al., 1988) that provides major ecosystem services (David and Handa, 2010), notably in agrosystems and grassland habitats where it is used as an ecological indicator. In the course of its evolutionary history, the common woodlouse had to adapt to terrestrial life. Consequently, it is still very sensitive to moisture and temperature variations, which can induce water loss (Smigel and Gibbs, 2008) and have major consequences in terms of distribution, behavior and survival (Hassall et al., 2018; Paris and Pitelka, 1962). Moreover, their movement capacity is low (i.e. several hundred meters during the entire lifetime at the best, Durand et al., 2019) to allow them to migrate so to avoid the stress imposed by the environment. Our knowledge and ability to measure woodlouse life history traits and the availability of molecular and cellular biomarkers of individual quality (Depeux et al., 2020a) make this species a highly relevant experimental model to study the influence of both temperature and moisture on life history and physiological traits.

In this study, we tested independently the effects of increased temperature (experiment 1) and of decreased moisture (experiment 2) on a selected set of key life history (i.e. growth, reproductive success, and survival) and physiological (i.e. immune cell parameters (cell viability, cell density and cell size) and β-galactosidase activity) traits in woodlouse. In experiment 1 (i.e. testing the effect of increased temperature), we compared individuals maintained at 20°C and 80% of moisture (i.e. the standard temperature and moisture laboratory conditions) to individuals exposed at 28°C (simulating a temperature increase of 8°C) still at 80% of moisture. In experiment 2 (i.e. testing the effect of decreased moisture), we compared animals in standard conditions to individuals exposed at 50% of moisture (simulating a moisture loss of 30%) still at 20°C. We hypothesized that a rise in temperature and a loss in moisture should be stressful and should induce changes in life history and physiological traits.

## MATERIALS & METHODS

### Biological Material – Routine Breeding

All specimens of *A. vulgare* used in this study descend from individuals sampled in Denmark (Helsingör) in 1982. Since then, animals were reared under laboratory conditions under the natural photoperiod of Poitiers (France) (46°35’N; 0°20’E), at 20°C and about 80-85% of moisture, in plastic boxes (length × width × height: 26.5 × 13.5 × 7.5 cm) containing humid loam, and fed ad libitum with carrot slices and dried linden leaves. Controlled breeding, for the maintenance of the lineage over years, is performed in individual boxes (diameter x height: 9,8cm x 4,9cm), with reproductive pairs selected from their pedigree to avoid inbreeding. One month after mating, offspring exit the female *marsupium* (i.e. female ventral pouch on which the eggs develop) (Suzuki and Ziegler, 2005). We transferred these offspring a few days after birth into a bigger box (length × width × height: 26.5 × 13.5 × 7.5 cm) with loam and food. After 3 months, once sexual characters have appeared but before sexual maturity, we placed young males and females in separate boxes (length × width × height: 26.5 × 13.5 × 7.5 cm) in laboratory conditions described above, enabling us to obtain virgin adults. For the maintenance of the lineage, about 40 crosses have been performed following this protocol each year. Each of the 40 broods provides at least one breeder for the next generation. The animals used in the experiments of this study came from this controlled lineage.

### Experimental Design

#### Experiment 1: effect of increased temperature on life history and physiological traits

The experiment 1 performed in January 2019 involved the comparison between two groups of animals aged of 7 months old maintained at different temperatures in two climatic chambers (Memmert HPP 256L with LED Light module cold white 6500K for HPP260 (15%) and Interior IP68 socket (for temperature restriction)) during two months after standard conditions of maintenance:

i. The “Control Temperature” group (CT) of animals maintained in standard conditions (i.e. at 20°C and 80% of moisture) in one of our two climatic chambers.
ii. The “High Temperature” (HT) group of animals exposed at 28°C (simulating increased temperature by 8°C (i.e. thermal stress condition)) and at 80% of moisture in the second climatic chamber.

Eight degrees (i.e. difference in temperature between the two groups) corresponds to a temperature increase close to daily variations observed in Poitiers during some summers, which could chronically induce stress. Moreover, we have observed the stressful effect of this temperature increase in a preliminary experiment in which we did not control the moisture variation (Depeux et al., 2019).

In each group, animals were fed *ad libitum* in 3 boxes (length × width × height: 26.5 × 13.5 × 7.5 cm; standard laboratory density conditions) of 30 females and 3 boxes of 30 males from 15 different clutches (i.e. all treatments included animals with the same genetic background (i.e. issued from 15 same clutches) to be comparable). For each condition, one box was used to monitor survival and growth (mass gain over time) of animals from the beginning to the end of the experiment, another was used to evaluate reproductive success and the last box served to quantify physiological traits (i.e. immune cell parameters: cell viability, cell density, and cell size) and β-galactosidase activity, see below). In this last box, the animals had to be sacrificed (see ‘Ethical statement’ section below) because of the protein extraction on nerve chains that was required to measure the β-galactosidase activity.

#### Experiment 2: effect of moisture loss on life history and physiological traits

The experiment 2 performed in January 2021 involved the comparison between two groups of 7 months old animals maintained under different conditions in our two climatic chambers during two months after standard conditions of maintenance:

i. The “Control Moisture” (CM) group of animals maintained in standard conditions (i.e. at 20°C and 80% of moisture) in one of our two climatic chambers
ii. The “Loss of Moisture” (LM) group of animals exposed at 50% of moisture (simulating a moisture loss of 30% (i.e. water stress condition)) and at 20°C in the second climatic chamber.

Similar to the experiment 1, in each group, animals were fed *ad libitum* in 3 boxes of 30 females and 3 boxes (length × width × height: 26.5 × 13.5 × 7.5 cm; standard laboratory density conditions) of 30 males from 15 different clutches (i.e. all boxes to compare included animals with the same genetic background (i.e. issued from 15 same clutches)). For each condition, one box was used to monitor individual survival and growth from the beginning to the end of the experiment, another was used to evaluate reproductive success and the last box served to quantify physiological traits (see below).

In our two experiments, we aimed to compare individuals of the same age because age negatively impacts both reproductive success (Depeux et al., 2020b) and physiological traits (Depeux et al., 2020a). Having initially only two climatic chambers, we had to perform our experiments 1 and 2 in different years (i.e. experiment 1 in 2019 and experiment 2 in 2021).

Thus, we systematically compared the effect of each stress against its own control condition group (i.e. CT for HT and CM for LM). Moreover, at the beginning of each experiment (1 and 2), we selected individuals of the same size and we checked, at the end of the experiments, potential statistical differences between the two control groups (CT and CM) on measures of life history and physiological traits (Supp. File1).

### Ethical statement

The Decree n°2003-768 from 01/08/2003 and the European Directive 2010/63/EU regulating animal research does not require any ethical evaluation prior to research on arthropods. However, we complied with the ethical 3R rules (Replace/Reduce/Refine). Even though it was impossible to replace the use of animals in our study, we reduced the number of used animals, optimizing this number to a minimum to ensure a reliable assessment of the effect of the different stressors on life history and physiological traits. Although individuals were obviously stressed during the experiments, we made sure that they were provided with optimal living conditions throughout the experiments. In addition, when the tissue sampling required the death of individuals (i.e. to measure physiological traits such as β-galactosidase activity), the animals to be euthanized were frozen before protein extractions to take into account animal welfare as much as possible.

### Life history traits

#### Survival and growth

One box of males and one box of females from each group (i.e. for the groups CT, HT, CM, LM) were used to monitor and compare changes of survival and body mass over time. All individuals in these boxes were monitored for 124 days (i.e. about 4 months). We sampled individuals at 14, 28, 42, 69, and 124 days (i.e. 5 sampling points per box) and assessed survivorship and change in body mass (in grams) of all surviving animals in each box (body mass was measured with a precision balance 650g|1mg Sartorius™ BCE653-1S Entris™ II Essential). Then, we compared these traits over time and between groups (CT vs. HT groups and CM vs. LM groups) to test independently the effect of temperature and moisture changes on these traits (see section on Statistical analyses). Due to regular moults, individual identification among the 30 animals sharing in given box cannot be performed, leading our measures to be average survival and growth instead of individual trajectories.

#### Reproductive success

At the end of the exposure to different conditions, one box of males and one box of females were collected from each group (i.e. for the groups CT, HT, CM, and LM). We formed 20 breeding pairs composed of one male and one female per group. Each breeding pair was placed in a box, at 20°C, with food provided *ad libitum* and in a photoperiod of 16:8 (L/D) stimulating the reproduction (Mocquard et al., 1989). We followed all these pairs for 5 months during which each clutch produced was recorded. At the end of this period, the ability to produce a clutch (i.e. the probability that one clutch or more is produced by a given pair) for the 80 pairs (i.e. 40 pairs for experiment 1 and 40 pairs for experiment 2) was compared independently between CT and HT groups and between CM and LM groups to test the effect of temperature and moisture changes on breeding success. As we created groups from similar clutches to have the same genetic background among boxes, we cannot exclude that some crosses were composed of related individuals although we expect this event to be rare. However, the probability of forming sibling pairs (8%) was similar among groups that were exposed either at 20°C vs. 28°C or at 80% vs. 50% of moisture.

### Physiological traits

At the end of the experimental treatments (i.e. after two months in our experimental conditions), one box of males and one box of females were taken from each group (i.e. for the groups CT, HT, CM, and LM) for measuring the level of our set of physiological traits (i.e. immune cells parameters and β-galactosidase activity) developed in Depeux et al. (2020a). These physiological traits were firstly described as senescence biomarkers because they allow predicting the amount of cellular senescence in different organisms and are strongly age-dependent in *A. vulgare* (i.e. older the individual, higher the decline of these biomarkers, Depeux et al. 2020a). We performed these measures on each remaining animals (Table 1) and we compared these metrics independently between CT and HT groups for experiment 1 and between CM and LM groups for experiment 2 (see section Statistical analyses).

**Table 1.**
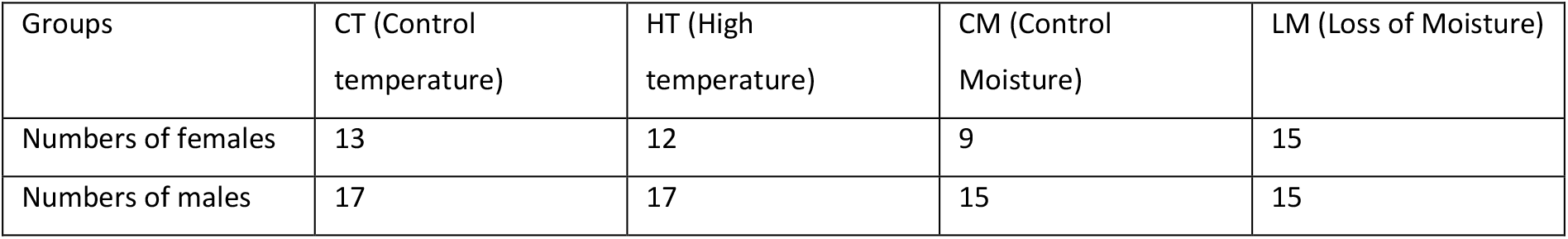
Numbers of individuals on which we measured quality biomarkers.

| Groups | CT (Control temperature) | HT (High temperature) | CM (Control Moisture) | LM (Loss of Moisture) |
| --- | --- | --- | --- | --- |
| Numbers of females | 13 | 12 | 9 | 15 |
| Numbers of males | 17 | 17 | 15 | 15 |

#### Immune cells

As immune cells are free-circulating, they can inform about a potential premature biological aging. When an individual *A. vulgare* ages, its immune cells decrease in density and viability while increasing in size (Depeux et al., 2020a). To measure these parameters, we collected 3μL of haemolymph per individual and placed it in 15μL of MAS-EDTA (EDTA 9 mM, Trisodium citrate 27 mM, NaCl 336 mM, Glucose 115 mM, pH 7, (Rodriguez et al., 1995)). We then added 6μL of Trypan Blue at 0.4% (Invitrogen) to discriminate live and dead cells. After, 10μL of this solution was put in Invitrogen Coutness® counting slide and put in an automated Cell Counter (Invitrogen) to quantify cell density (measured as the number of cells per mL of haemolymph), viability (measured as the proportion of live cells) as well as cell size (in μm). These three parameters of the immune cells are physiological traits that were found to be reliable biomarkers of cellular senescence in *A. vulgare* (Depeux et al., 2020a).

#### β-galactosidase activity

The β-galactosidase activity is a physiological trait commonly used as a marker of cellular senescence (Lee et al., 2006). Its indirect activity in regards to the process of cellular senescence increases with age in *A. vulgare* (Depeux et al., 2020a). To measure this enzymatic activity, we dissected and removed the nerve cord of each individual after having collected haemolymph for assessing the immune parameters. We put individual nerve cords in 300μL of Lyse Buffer 1X (CHAPS 5 mM, Citric acid 40 mM, Sodium Phosphate 40 mM, Benzamidine 0.5 mM, PMSF 0.25 mM, pH = 6) (Gary and Kindell, 2005). We centrifuged the sample at 15 000g for 30 minutes at 4°C and then we collected and kept the supernatant at −80°C. We quantified the protein concentration thanks to the BCA Assay and we homogenized all samples at the 0.1mg/mL protein concentration. Then, 100μL of these protein extracts were added to 100μL of reactive 4-methylumbelliferyl-D-galactopyranoside (MUG) solution. The synthesis of the fluorescent 4-methylumbelliferone (4-MU), the result of the contact of MUG reactive with β-galactosidase, was measured using the multimode microplate reader Mithras LB940 133 HTS III, Berthold; excitation filter: 120 nm, emission filter 460 nm, for 120 minutes. We included two technical replicates for each sample to obtain the measures.

### Statistical analyses

All statistical analyses were performed using the software R 4.2.1 (R core Team 2022). The effects of the stress condition (control vs. high temperature, or control vs. low moisture) on life history and physiological traits were tested using the following models. (*i*) Life history traits. For the survival data, Cox proportional hazard models were fitted with stress condition, sex and their interaction term as fixed variables, using the ‘survival’ package (Therneau, 2022). For the growth data, the body mass was modelled using linear models with Gaussian distribution with stress condition, sex, time (i.e. time after placing in climatic chamber, in days) and their two-by-two interaction term as fixed variables. The female reproductive success was modelled as binary data (*presence of at least one clutch or absence of clutch*) using linear regression with a binomial distribution, with stress condition as fixed variables. (*ii*) Physiological traits. The cell density, cell size, cell viability, and the β-galactosidase activity were modelled using linear models with Gaussian distribution with stress condition, sex, and their interaction term as fixed variables.

We proceeded to model selection starting with full (saturated) model. We ranked all nested models according to their AICc using ‘MuMIn’ package (Barton, 2022). We selected the most parsimonious models among the top ranked models (ΔAICc < 2) (Galipaud et al., 2017). The tables summarizing the model selection procedure was presented in Supp. File2. To represent the effect of the two environmental stresses in each variable, we presented our results with indices of size effect. The effect of each stress on each measure of life history traits and biomarkers of individual quality were measured using the standardized slopes (Schielzeth et al 2010) and their SE calculated by rescaling the variable of the selected model. For survival data, the effect size was the hazard ratio, calculated as the exponential of the regression parameter (Collett 2003). When the selected model did not include the effect of the stress, we took the model with the variable stress as fixed factor to obtain a size effect as done in Depeux et al. 2020a.

### Data, script and code availability

All datasets and source code are available as electronic supplementary materials on public repository: https://doi.org/10.5281/zenodo.7496837

## RESULTS

As said previously, we checked, at the end of the experiments, potential statistical differences between the two control groups (CT and CM) on measures of life history and physiological traits (Supp. File1). Although β-galactosidase activity and cell density were higher in the CT group than in the CM group (Supp. File1), we observed the same dynamics in these measures in the face of their stressful condition (HT and LM, respectively) (see Results part). The body mass at day 14 was higher in the CM group than in the CT group (Supp. File1). Whatever the differences observed between the two control groups (CT group used in 2019 and CM group used in 2021), we compared the effect of each stress (HT in 2019 and LM in 2021, respectively) against its own control group (CT group in 2019 and CM group in 2021, respectively) for testing the effect of each stress.

### Life history traits

Survival was not impacted by an increased temperature (χ^2^_1_=2.16, P=0.14, Fig.1A, Supp. File2 Table S2a, Supp. File3-1.A.1) although mortality risk was almost twice as lower at low compared to high temperature. The hazard ratio was 1.78, with a 95%CI including 1 [0.81; 3.88]. By contrast, individuals exposed to a water stress had a 2.5 times higher mortality risk (χ^2^_1_=4.54, P=0.03, Fig. 1B, Supp. File2 Table S2b, Supp. File3-1.A.2). The hazard ratio was 2.69, with a 95%CI excluding 1 [1.03; 7.01]. As a result, 90% of individuals placed at control moisture were still alive at the end of the follow-up, whereas only 75% of individuals placed in water stress condition survived at the end of the experiment.

**Figure 1.**
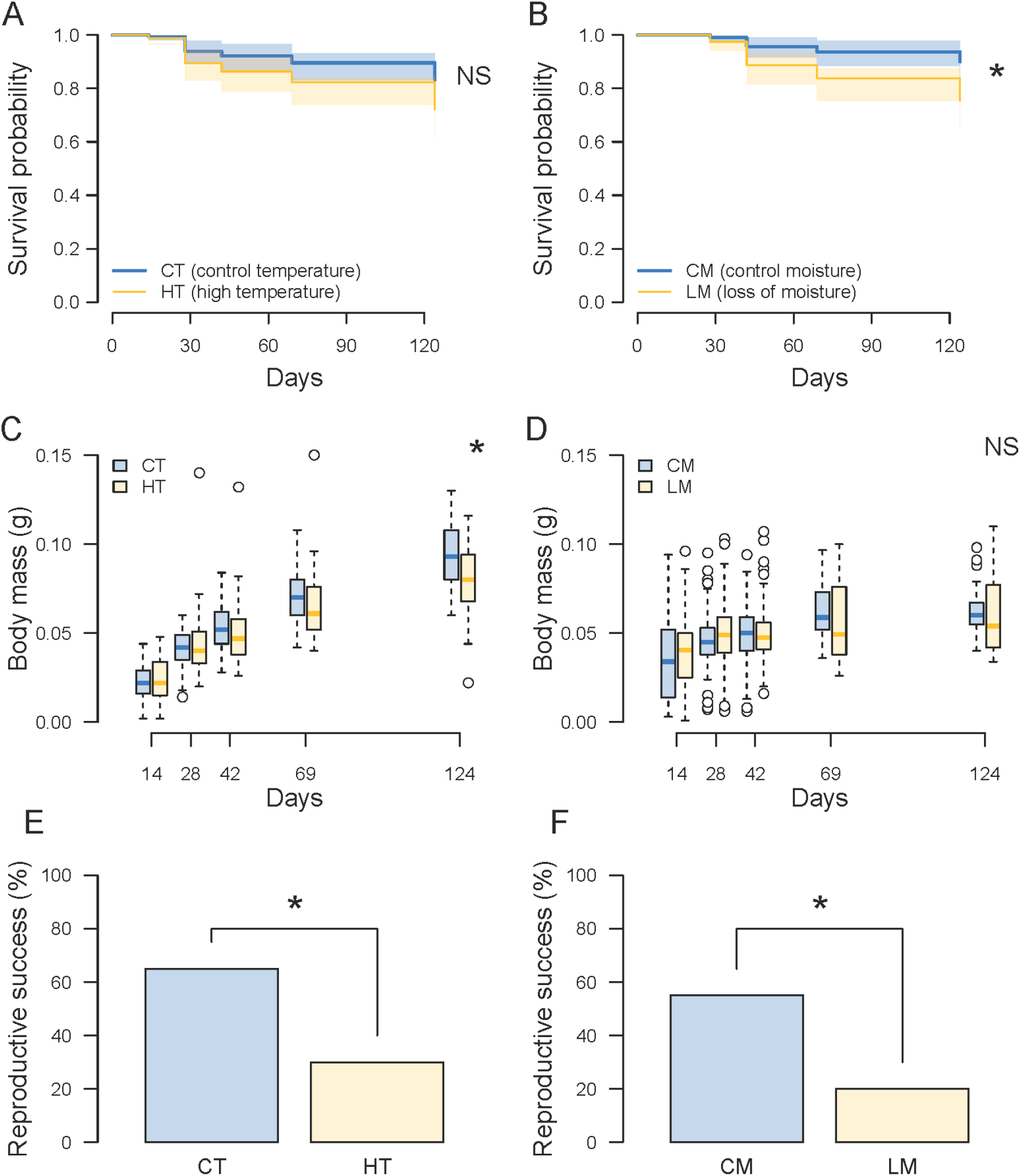
Effect of the two environmental stressors (Temperature (A, C and E) and Moisture (B, D and F) on Survival (A and B), Body mass (C and D) and Reproductive success (E and F). Blue colour: control groups, orange colour: stress groups. CT: Control Temperature (20°C), HT: High Temperature (28°C), CM: Control Moisture (80%), LM: Loss of Moisture (50%). NS: No significant; * p<0.05.

In both thermal and water stresses, body mass increased during the entire experiment duration (Fig. 1C and 1D, Supp. File3-1.B.1 and 1.B.2), as expected in an indeterminate grower as *A. vulgare*, but there was no detectable interaction between day and sex (Supp. File2 Table S2c and Table S2d). For the temperature experiment, interactions between sex and stress (F_1,528_= 6.90, P=0.0088, Supp. File3), and between day and thermal stress (F_1,528_=14.6, P=0.00015, Supp. File2 Table S2c) showed up, illustrating the impact of an increasing temperature on growth. By contrast, in the moisture experiment, the body mass was not affected by the sex (F_1,522_=0.35, P=0.55), the water stress (F_1,522_=0.31, P=0.58), or any interaction between the variables (all P > 0.10).

The reproductive success markedly decreased in both experiments for the stressful condition: a fourfold increase of reproductive failure in presence of thermal stress (χ^2^_1_=5.02, p=0.025, Odd-ratio=0.23, 95%CI=[0.057;0.83], Fig. 1E, Supp. File2 Table S2e, Supp. File3-1.C.1), and a fivefold increase of reproductive failure in presence of water stress (χ^2^_1_=5.38, p=0.02, Odd-ratio=0.20, 95%CI=[0.045;0.79], Fig. 1F, Supp. File2 Table S2f, Supp. File 3-1.C.2). In both cases, it corresponds to halving the reproductive success in the stress groups (water stress: 55% in the control group vs. 20% in the stressed group; thermal stress: 65% in the control group vs. 30% in the stressed group).

### Physiological traits

#### Immune cells

Immune cell viability was not affected by the thermal stress (F_1,56_=0.92, p=0.34, standardized slope ß=-0.25; 95%CI=[-0.79;0.27], Fig. 2A, Supp. File2 Table S2g and Supp. File3-2.A.1) but decreased during the water stress (F_1,50_=4.17, p=0.046, standardized slope β=0.55; 95%CI=[0.01;1.09], Fig. 2B, Supp. File2 Table S2h and Supp. File3-2.A.2).

**Figure 2.**
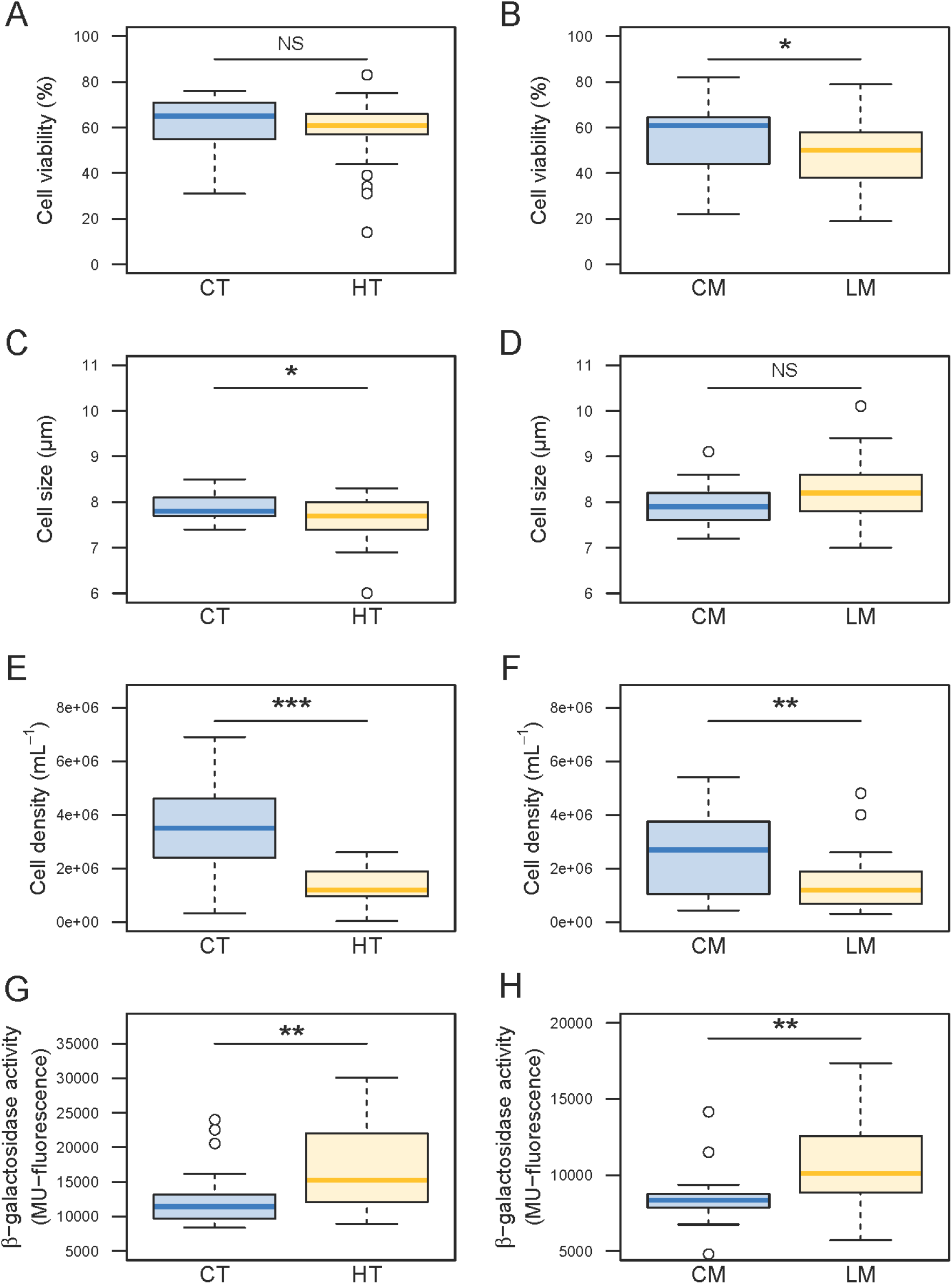
Effect of the two environmental stressors (Temperature (A, C, E, and G) and Moisture (B, D, F and H) on immune cell viability (A and B), immune cell size (C and D), immune cell density (E and F) and β-galactosidase activity (G and H). Blue colour: control groups, orange colour: stress groups. CT: Control Temperature (20°C), HT: High Temperature (28°C), CM: Control Moisture (80%), LM: Loss of Moisture (50%). NS: No significant; * p<0.05, ** p<0.01, *** p<0.001.

Immune cell size decreased during the thermal stress (F_1,55_=5.72, p=0.02, standardized slope β=-0.60; 95%CI=[-0.1;-1.1], Fig. 2C, Supp. File2 Table S2i and Supp. File 3-2.B.1) but not under the water stress (F_1,50_=3.79, p=0.057, standardized slope β=-0.55; 95%CI=[-1.07;0.02], Fig. 2D, Supp. File2 Table S2j and Supp. File3-2.B.2).

Immune cell density decreased during the thermal stress (F_1,56_= 38.2, P<0.001, standardized slope β=-1.26; 95%CI=[-1.67;-0.85], Fig. 2E, Supp. File2 Table S2k and Supp. File3-2.C.1)) and the water stress (F_1,50_=7.64, p=0.008, standardized slope β=0.72; 95%CI=[0.19;1.25], Fig. 2F, Supp. File2 Table S2l and Supp. File3-2.C.2).

#### β-galactosidase activity

The *β*-galactosidase activity increased with the thermal stress (F_1,54_=11.32, P=0.0014, standardized slope β=0.82; 95%CI=[0.33;1.32], Fig.2G, Supp. File2 Table S2m, and Supp. File3-2.D.1), but also with the water stress (F_1,50_=10.50, P=0.002, standardized slope β=-0.83; 95%CI=[-1.31;-0.32], Fig. 2H, Supp. File2 Table S2n and Supp. File3-2.D.2).

## DISCUSSION

Our results highlight that life history traits were negatively impacted by the two environmental stressors (thermal and water stresses) considered in this study. Moreover, the detrimental effects of these stressors on our set of biomarkers of individual quality are consistent with an overall premature ageing of stressed animals compared to unstressed ones. To briefly summarize, an increase in temperature (thermal stress) negatively affects both the body mass trajectory over time and the reproductive success of individuals. A decrease in moisture (water stress) resulted in a decrease of both survival and reproductive success. Concerning our physiological traits: (1) the density of immune cells decreases under both stresses, (2) immune cell size decreases under thermal stress, but is not impacted under water stress, (3) the viability of the cells decreases under water stress (but not under thermal stress) and (4) finally, the β-galactosidase activity increases for the two stressed groups. In this context, our results globally support marked negative effects of thermal and water stresses on woodlouse performance, with some minor differences between the two stressors in their effects on life history and physiological traits.

About the life history traits, if the thermal stress has no detectable effect on survival in *A. vulgare*, contrary to what has been previously reported in arthropods studied so far such as *Antestiopsis thunbergii, Calliphora stygia* and *Margaritifera margaritifera* (Azrag et al., 2017; Hassall et al., 2017; Kelly et al., 2013), this stressor leads to a slowdown in woodlouse growth, in line to what has been reported in three other isopods (Angilletta et al., 2004). In parallel, the water stress leads to a decrease in reproductive success, as previously reported in females of *Antestiopsis thunbergii* (Azrag et al., 2017). In *A. vulgare*, individual body size is positively correlated with fecundity (Durand et al., 2018; Lawlor, 1976), meaning that the slowdown in growth could explain, at least partly, the decrease in reproductive success for stressed animals compared to non-stressed ones. Concerning the water stress, if the loss of moisture has no detectable effect on woodlouse growth, it causes a decrease in both survival and reproductive success. These findings suggest a high cost of drought on individual fitness in *A. vulgare*.

About the physiological traits, our results of the thermal stress experiment show that although cell density is negatively impacted by increased temperature, cell viability is not affected. Moreover, contrary to the expectation when individuals are senescent, cell size decreases instead of increasing. This last result supports our previous finding that cells decrease in size when the temperature raises (without controlling moisture level, Depeux et al., 2019). That smaller cells are associated with increased cell renewal in stressed animals might explain this pattern. On the other hand, the increase of β-galactosidase activity seems to indicate premature ageing (and thus a decrease in quality) in individuals exposed to thermal stress. Concerning the water stress, the biomarkers of individual quality indicate a decrease in cell density and viability, associated with an increase in β-galactosidase activity, which suggests an acceleration of biological ageing in the individuals exposed to a water stress (Depeux et al., 2020a).

We reported a global negative effect of the thermal stress in *A. vulgare* in our study, but our results seem to show an even higher and clearer effect of the water stress on both life history traits and biomarkers of individual quality. Although the woodlouse has become terrestrial for a long time, the individuals of that species are still dependent on and require a substantial water supply (Smigel and Gibbs, 2008). Thus, behaviours like aggregation that allow individuals to resist to desiccation have been set up and thereby to maintain the rate of moisture required for survival (Broly et al., 2013; Smigel and Gibbs, 2008). This can explain the strong effect of water stress in our study. Under natural conditions, increase in temperature and loss of moisture generally positively covary, leading to even higher negative consequences on the woodlouse performance. Further work will be required to test the influence of more extreme and maybe more realistic conditions by simultaneously increasing temperature and decreasing moisture on life history and physiological traits. A study in the wild comparing life history and physiological traits on *A. vulgare* collected across areas with different temperature and drought gradient would also allow a better assessment of the combined effects of these stressors.

Unlike what happens in nature, our experimental study on a laboratory line of woodlouse allowed us to test the effect of the thermal and water stresses while controlling for potentially confounding factors such as individual age. Indeed, it is highly challenging to control for individual age in the wild. In this context, the use of our controlled laboratory line on which we developed our physiological markers allowed us to account for the exact age of the animals (which is itself linked to life history and physiological traits (Depeux et al., 2020b)) and for their genetic origin. We compared groups of the same origin (and our controlled crosses guarantee the genetic diversity of our line) and of the same age. This allows us to limit confounding effects as much as possible and to quantify the effects of the two tested stressors independently.

Thanks to our experimental design that allowed us to test independently the influence of stressors that organisms are likely to face in the wild, we showed that thermal and water stress do not have the same impact. Although simulations based on mathematical models have predicted that both temperature and drought changes overall affect arthropods, experimental approaches such as reported in this work are required to quantify reliably the influence of changing conditions on life history and physiological traits (Johnson et al. 2010). Drought can have serious physiological consequences on invertebrates, involving e.g. protein denaturation and undesirable macromolecular interactions (Sano et al., 1999; Tang and Pikal, 2005) or oxidative damage (Lopez-Martinez et al., 2008), which are known to be associated with cellular senescence (Gilca et al., 2007) and thus in the decreased performance observed in stressed organisms. Due to the role of arthropods in services to many ecosystems (e.g. biochemical balance of ecosystems, agriculture, pest management...), and in the context of global warming, it is crucial to understand the effects of temperature and moisture changes on these organisms (Santos et al., 2021). As temperature increase is not the only environmental change expected to take place in the coming years, it is of paramount importance to assess also the impact of other stressors. Although many predictive models have been proposed so far, getting more accurate information on the expected responses of organisms facing with different kinds of stress would provide the required information to test these model predictions.

To conclude, *A. vulgare* is an important actor that delivers ecosystem services in many ecosystems because it actively impacts soil fertility (Souty-Grosset and Faberi, 2018) and it is also used as an ecological indicator of grassland habitats (Paoletti and Hassall, 1999; Souty-Grosset et al., 2005). This detritivorous species facilitates decomposition processes and nutrient cycling on which agricultural productivity and sustainability depend (Bredon et al., 2018; Paoletti and Hassall, 1999), and plays thereby a key role in ecosystem services (David and Handa, 2010). Extending knowledge in the response of soil biodiversity facing current global changes could promote sustainability by helping to the development of new tools and strategies for more efficient management of soils and associated crops, through more effective and targeted recolonisation and/or restoration of soil biodiversity. Also, to better understand what the future of the animal communities in the current context of global warming will be, it is necessary to perform studies on models presenting particular ecological requirements, such as woodlouse.

## Supplementary files

All supplementary files are available on public repository: https://www.biorxiv.org/content/10.1101/2022.09.26.509512v1.supplementary-material Supp. File1 Comparison of the two control groups

Supp. File2 Model selection

Supp. File3 Graphical representations of results per sex

## ACKNOWLEDGMENTS

We would like to thank Alexandra Lafitte for technical assistance, and two anonymous reviewers and the recommender of PCI Ecology for their constructive comments.

## Supplementary File 1: Comparison of the two control groups

**Table 1:**
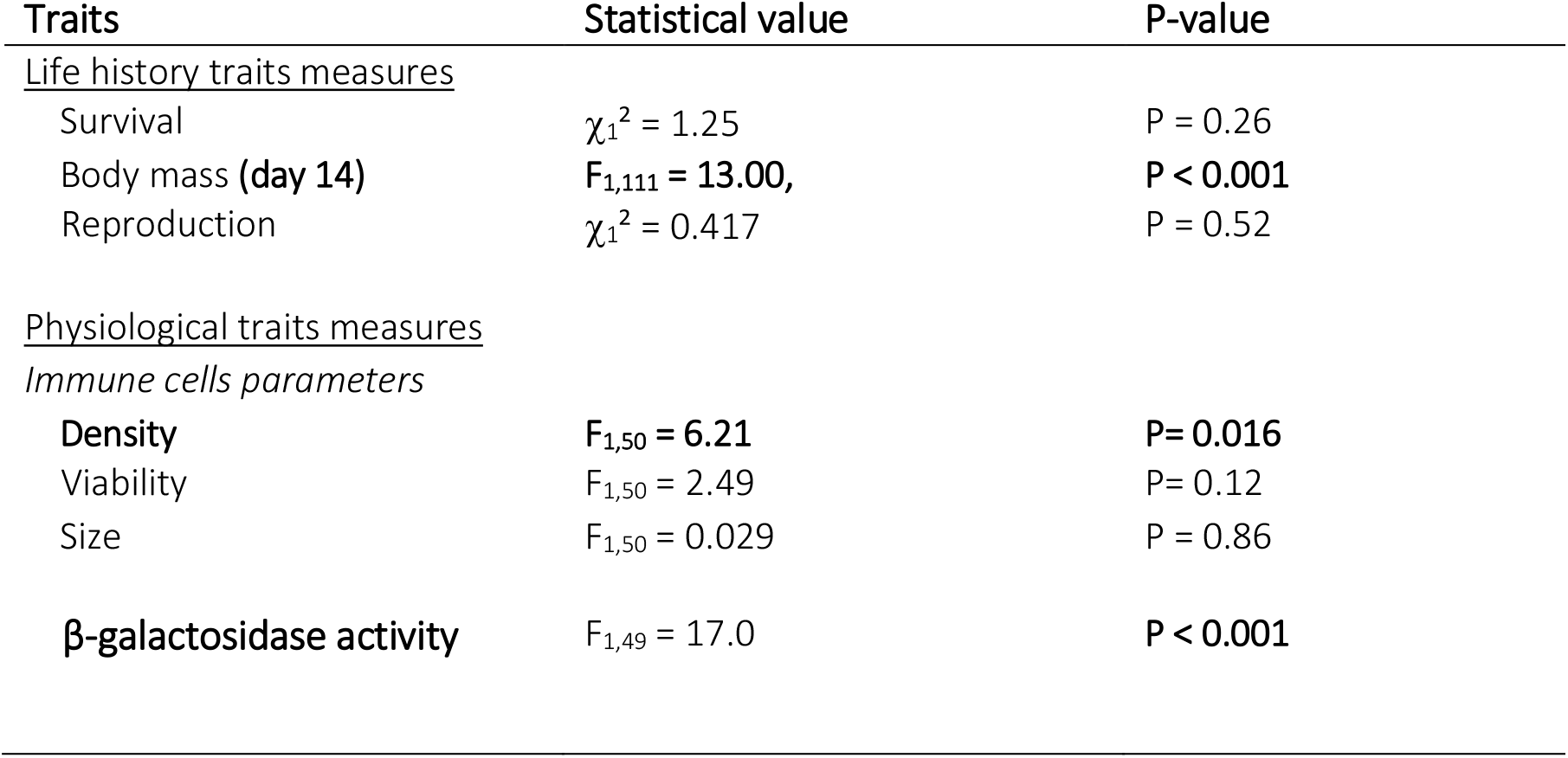
Comparison between the two control groups (CT (Control Temperature) and CM (Control Moisture)) of the two experiments for each tested variable (in bold the variables with significant statistical differences with graphical associated figures (Fig. 1, Fig. 2 and Fig. 3))

**Figure 1:**
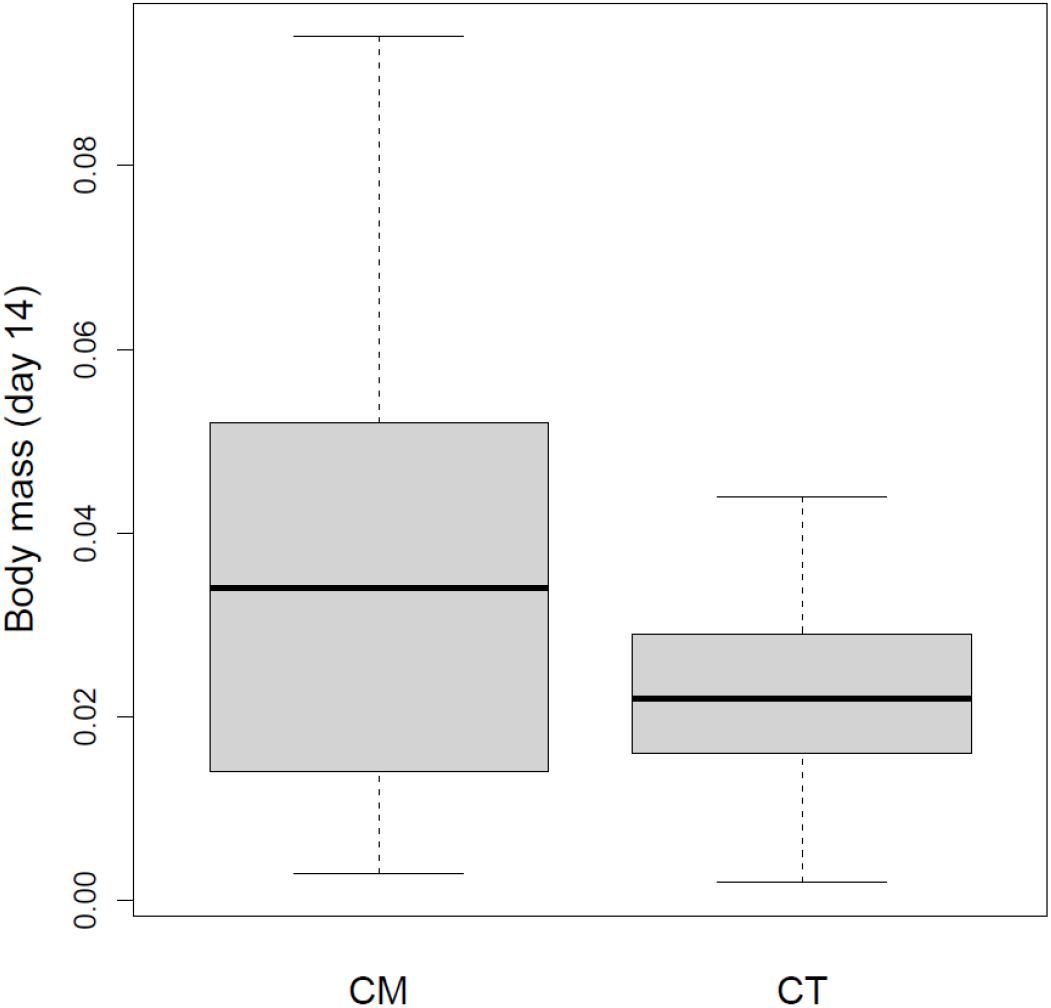
Body mass comparison between the two control groups (CT (Control Temperature) and CM (Control Moisture)) P-value < 0.001

**Figure 2:**
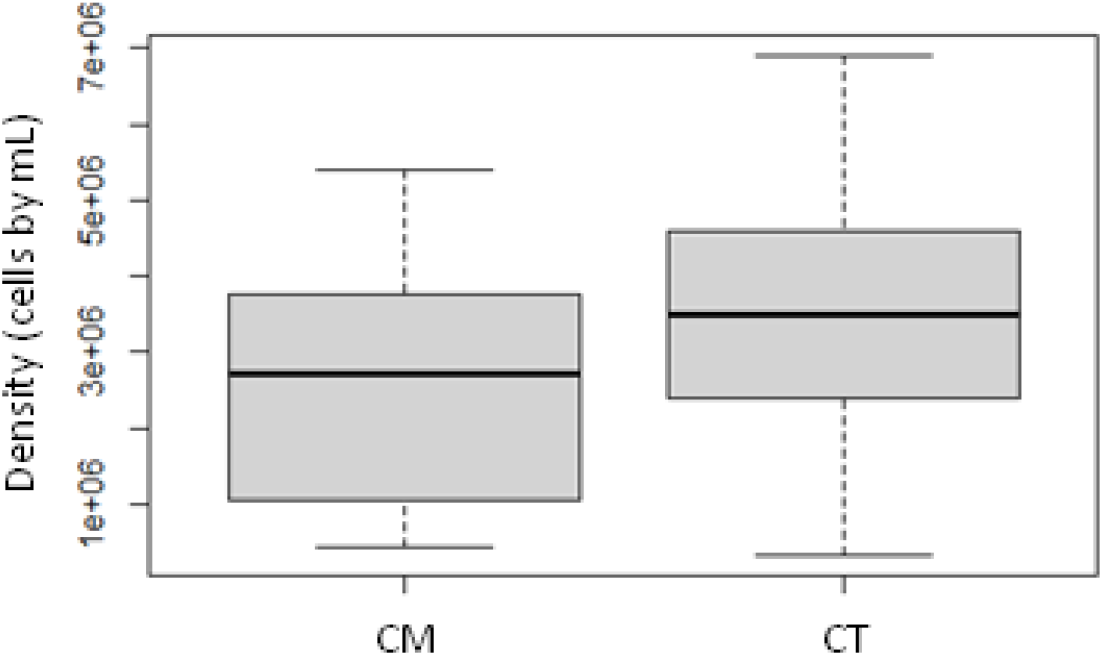
Immune cells density comparison between the two control groups (CT(Control Temperature) and CM (Control Moisture)) P-value=0.02

**Figure 3:**
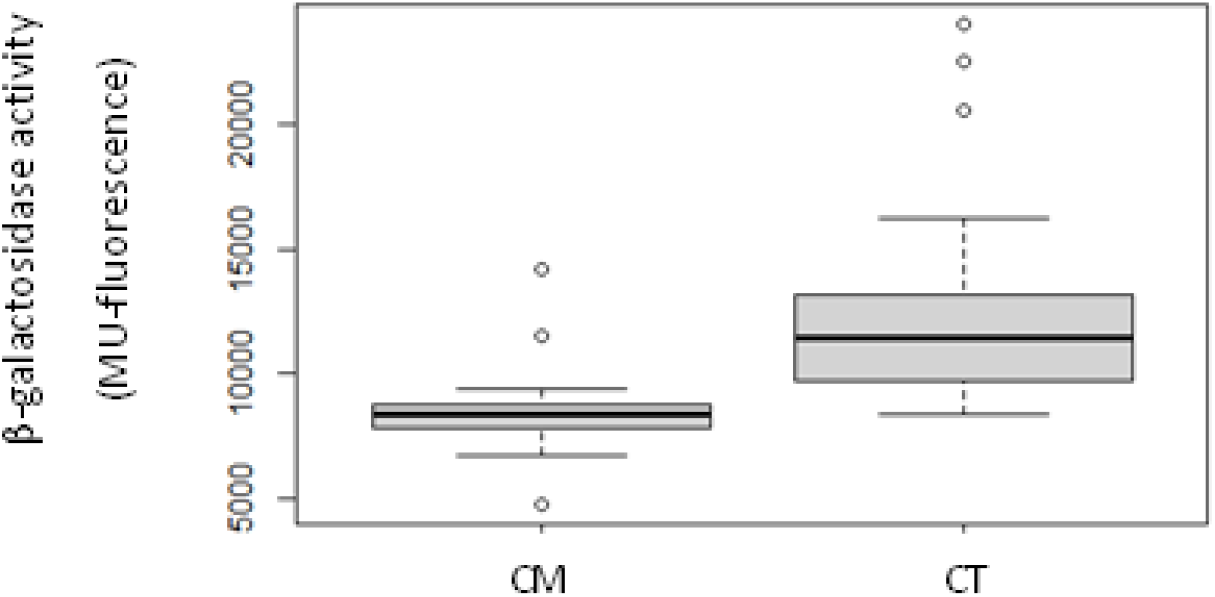
β-galactosidase activity comparison between the two control groups (CT (Control Temperature) and CM (Control Moisture)) P-value<0.001

## Supplementary File 2: Model selection

### Life history trait

#### Survival

**Table S2a.**
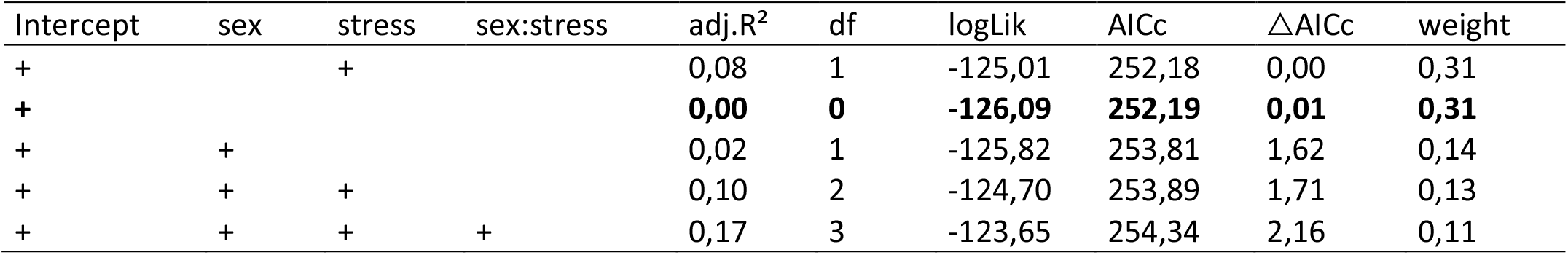
Effect of the thermal stress condition and sex on the survival. For each model, we reported intercept of the regression, adjusted R^2^ (adj.R^2^), degree of freedom (df), Log likelihood (LogLik) values, Akaike information criteria values with a correction for small sample sizes (AICc), change in AICc (ΔAICc) from the best model, and model weight. The presence of the categorial variable (sex, stress condition, and their interaction term sex:stress) in the model is indicated by a “+” symbol.

| Intercept | sex | stress | sex:stress | adj.$R^2$ | df | logLik | AICc | ΔAICc | weight |
| --- | --- | --- | --- | --- | --- | --- | --- | --- | --- |
| + |  | + |  | 0,08 | 1 | -125,01 | 252,18 | 0,00 | 0,31 |
| + |  |  |  | **0,00** | **0** | **-126,09** | **252,19** | **0,01** | **0,31** |
| + | + |  |  | 0,02 | 1 | -125,82 | 253,81 | 1,62 | 0,14 |
| + | + | + |  | 0,10 | 2 | -124,70 | 253,89 | 1,71 | 0,13 |
| + | + | + | + | 0,17 | 3 | -123,65 | 254,34 | 2,16 | 0,11 |

**Table S2b.**
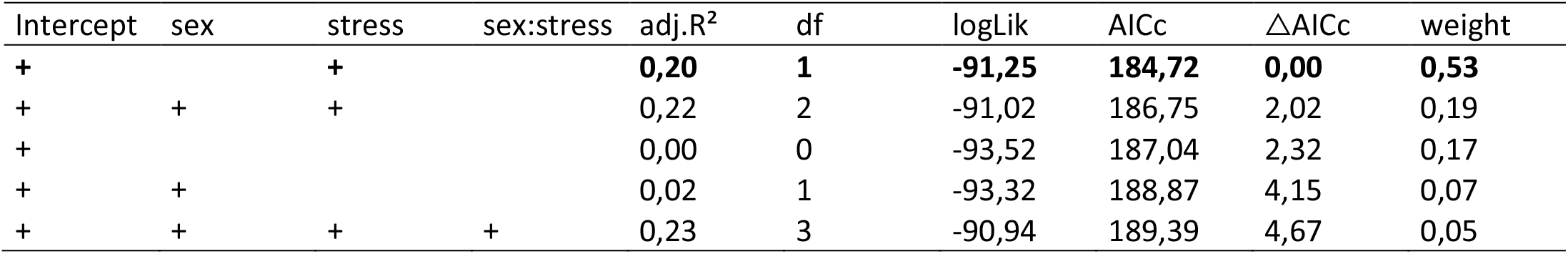
Effect of the water stress condition and sex on the survival. For each model, we reported intercept of the regression, adjusted R^2^ (adj.R^2^), degree of freedom (df), Log likelihood (LogLik) values, Akaike information criteria values with a correction for small sample sizes (AICc), change in AICc (ΔAICc) from the best model, and model weight. The presence of the categorial variable (sex, stress condition, and their interaction term sex:stress) in the model is indicated by a “+” symbol. The most parsimonious model is highlighted in bold font.

| Intercept | sex | stress | sex:stress | adj.$R^2$ | df | logLik | AICc | ΔAICc | weight |
| --- | --- | --- | --- | --- | --- | --- | --- | --- | --- |
| + |  | + |  | **0,20** | **1** | **-91,25** | **184,72** | **0,00** | **0,53** |
| + | + | + |  | 0,22 | 2 | -91,02 | 186,75 | 2,02 | 0,19 |
| + |  |  |  | 0,00 | 0 | -93,52 | 187,04 | 2,32 | 0,17 |
| + | + |  |  | 0,02 | 1 | -93,32 | 188,87 | 4,15 | 0,07 |
| + | + | + | + | 0,23 | 3 | -90,94 | 189,39 | 4,67 | 0,05 |

#### Body mass

**Table S2c.**
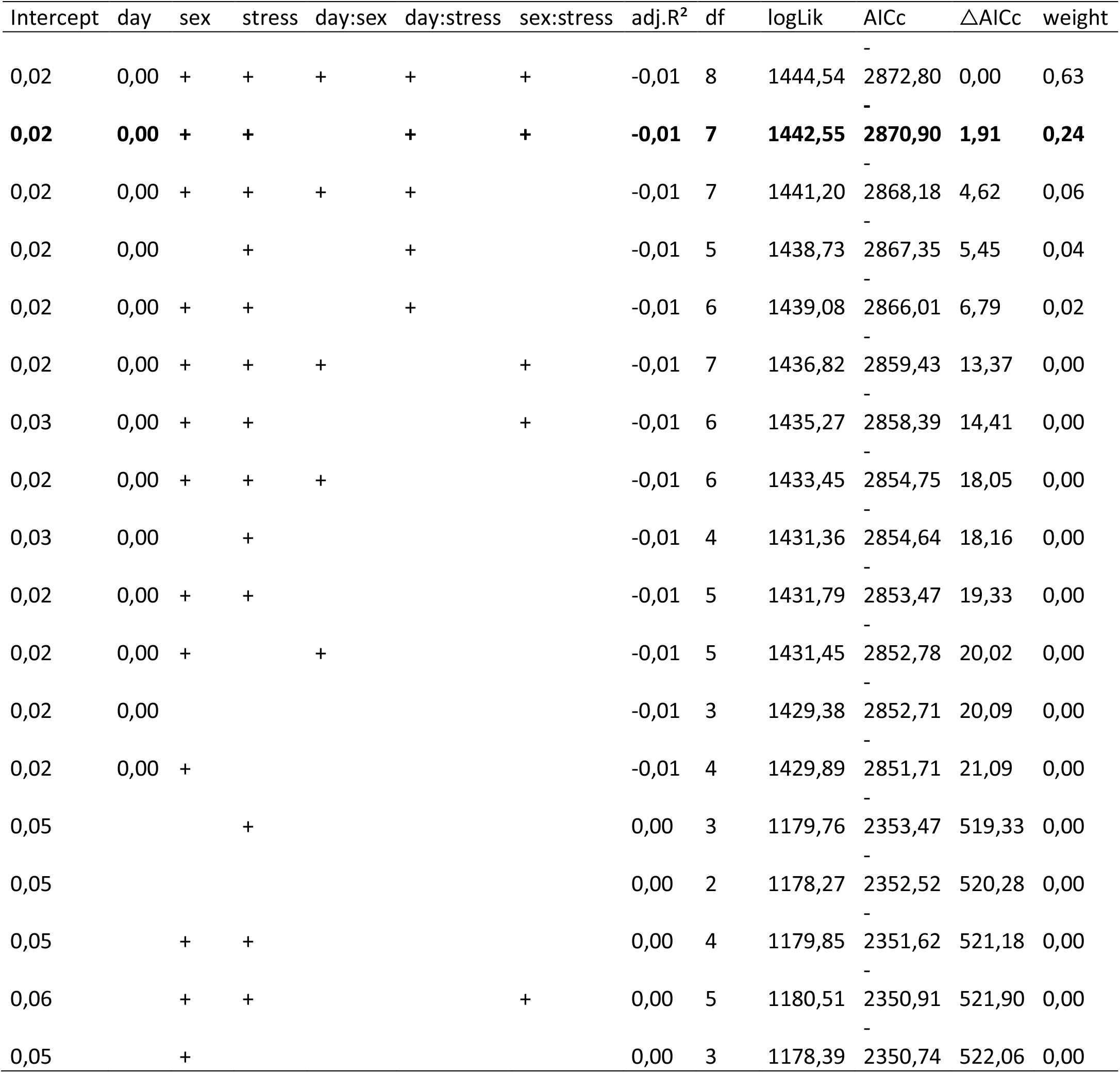
Effect of the thermal stress condition, sex and day on body mass. For each model, we reported intercept of the regression, adjusted R^2^ (adj.R^2^), degree of freedom (df), Log likelihood (LogLik) values, Akaike information criteria values with a correction for small sample sizes (AICc), change in AICc (ΔAICc) from the best model, and model weight. The presence of the categorial variable (sex, stress condition, and their two-by-two interaction terms) in the model is indicated by a “+” symbol. The regression parameter is only given for the corresponding continuous variable (day) when this variable is present in the model. The most parsimonious model is highlighted in bold font.

**Table S2d.**
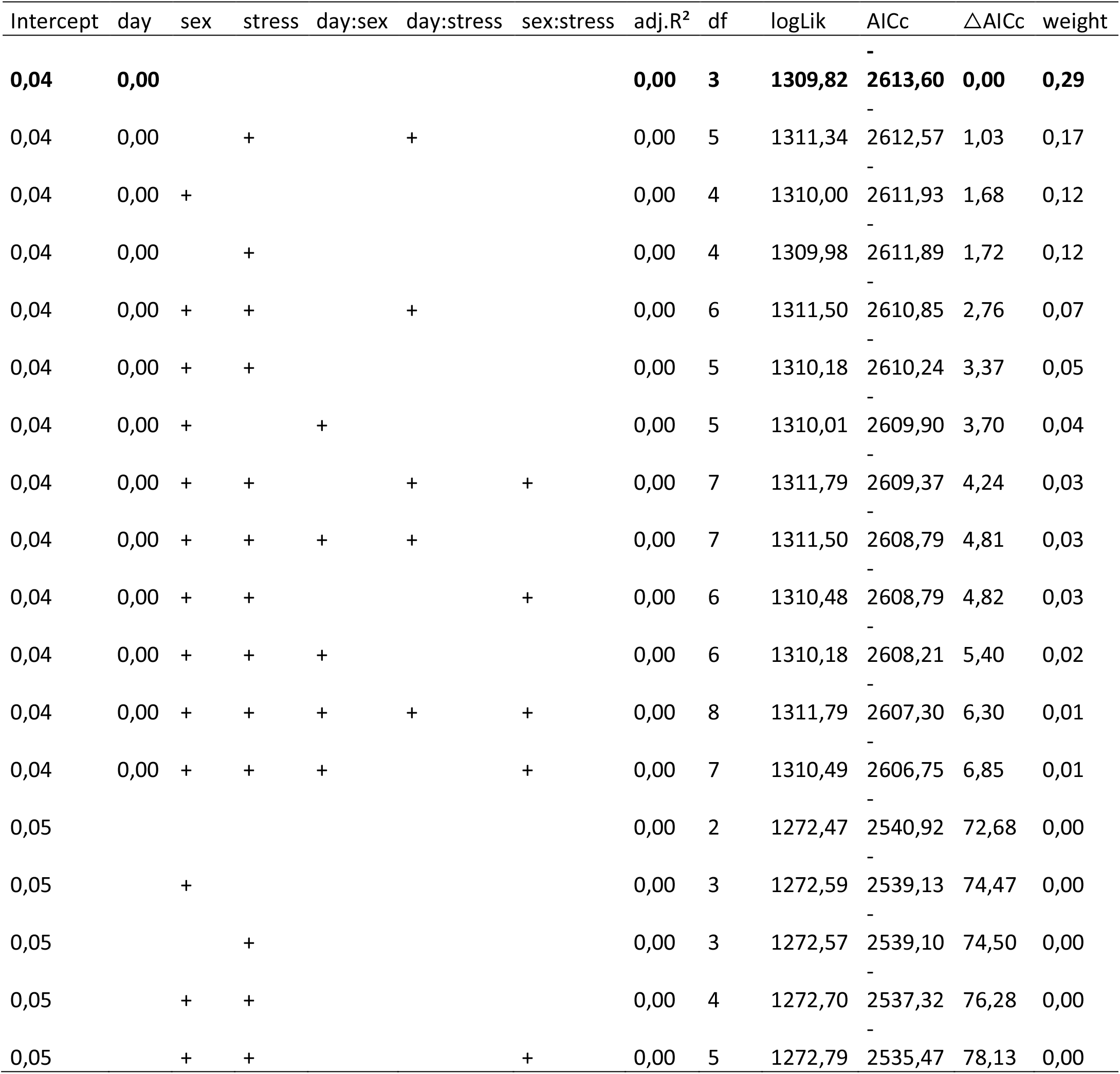
Effect of the water stress condition, sex and day on body mass. For each model, we reported intercept of the regression, adjusted R^2^ (adj.R^2^), degree of freedom (df), Log likelihood (LogLik) values, Akaike information criteria values with a correction for small sample sizes (AICc), change in AICc (ΔAICc) from the best model, and model weight. The presence of the categorial variable (sex, stress condition, and their two-by-two interaction terms) in the model is indicated by a “+” symbol. The regression parameter is only given for the corresponding continuous variable (day) when this variable is present in the model. The most parsimonious model is highlighted in bold font.

#### Reproductive success

**Table S2e.**
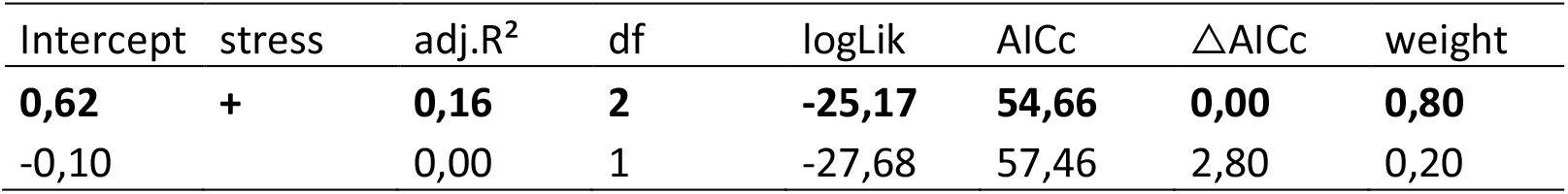
Effect of the thermal stress condition on the reproductive success. For each model, we reported intercept of the regression, adjusted R^2^ (adj.R^2^), degree of freedom (df), Log likelihood (LogLik) values, Akaike information criteria values with a correction for small sample sizes (AICc), change in AICc (ΔAICc) from the best model, and model weight. The presence of the categorial variable (stress condition) in the model is indicated by a “+” symbol. The value of regression parameter is only given for the intercept. The most parsimonious model is highlighted in bold font.

| Intercept | stress | adj.$R^2$ | df | logLik | AICc | ΔAICc | weight |
| --- | --- | --- | --- | --- | --- | --- | --- |
| **0,62** | **+** | **0,16** | **2** | **-25,17** | **54,66** | **0,00** | **0,80** |
| -0,10 |  | 0,00 | 1 | -27,68 | 57,46 | 2,80 | 0,20 |

**Table S2f.** Effect of the water stress condition on the reproductive success. For each model, we reported intercept of the regression, adjusted R^2^ (adj.R^2^), degree of freedom (df), Log likelihood (LogLik) values, Akaike information criteria values with a correction for small sample sizes (AICc), change in AICc (ΔAICc) from the best model, and model weight. The presence of the categorial variable (stress condition) in the model is indicated by a “+” symbol. The value of regression parameter is only given for the intercept. The most parsimonious model is highlighted in bold font.

| Intercept | stress | adj.$R^2$ | df | logLik | AICc | ΔAICc | weight |
| --- | --- | --- | --- | --- | --- | --- | --- |
| **-1,39** | **+** | **0,17** | **2** | **-23,77** | **51,87** | **0,00** | **0,83** |
| -0,51 |  | 0,00 | 1 | -26,46 | 55,03 | 3,16 | 0,17 |

### Individual physiological traits

#### Immune cell viability

**Table S2f.**
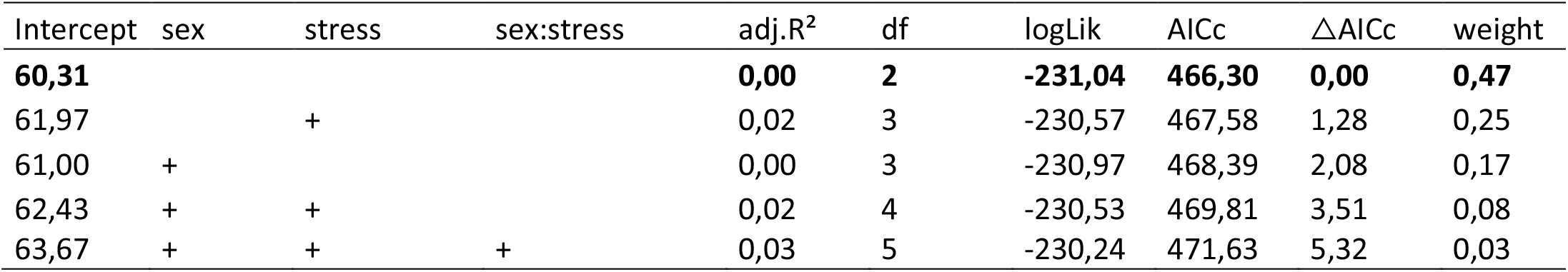
Effect of the thermal stress condition and sex on immune cell viability. For each model, we reported intercept of the regression, adjusted R^2^ (adj.R^2^), degree of freedom (df), Log likelihood (LogLik) values, Akaike information criteria values with a correction for small sample sizes (AICc), change in AICc (ΔAICc) from the best model, and model weight. The presence of the categorial variable (sex, stress condition, and their interaction term sex:stress) in the model is indicated by a “+” symbol. The value of regression parameter is only given for the intercept. The most parsimonious model is highlighted in bold font.

**Table S2h.** Effect of the water stress condition and sex on immune cell viability. For each model, we reported intercept of the regression, adjusted R^2^ (adj.R^2^), degree of freedom (df), Log likelihood (LogLik) values, Akaike information criteria values with a correction for small sample sizes (AICc), change in AICc (ΔAICc) from the best model, and model weight. The presence of the categorial variable (sex, stress condition, and their interaction term sex:stress) in the model is indicated by a “+” symbol. The value of regression parameter is only given for the intercept. The most parsimonious model is highlighted in bold font

| Intercept | sex | stress | sex:stress | adj.$R^2$ | df | logLik | AICc | ΔAICc | weight |
| --- | --- | --- | --- | --- | --- | --- | --- | --- | --- |
| 51,82 | + | + |  | 0,15 | 4 | -212,23 | 433,32 | 0,00 | 0,48 |
| **47,45** |  | + |  | **0,08** | **3** | **-214,36** | **435,21** | **1,90** | **0,19** |
| 52,29 | + | + | + | 0,15 | 5 | -212,20 | 435,70 | 2,39 | 0,15 |
| 55,52 | + |  |  | 0,06 | 3 | -214,86 | 436,23 | 2,91 | 0,11 |
| 51,29 |  |  |  | 0,00 | 2 | -216,44 | 437,13 | 3,81 | 0,07 |

#### Immune cell size

**Table S2i.** Effect of the thermal stress condition and sex on immune cell size. For each model, we reported intercept of the regression, adjusted R^2^ (adj.R^2^), degree of freedom (df), Log likelihood (LogLik) values, Akaike information criteria values with a correction for small sample sizes (AICc), change in AICc (ΔAICc) from the best model, and model weight. The presence of the categorial variable (sex, stress condition, and their interaction term sex:stress) in the model is indicated by a “+” symbol. The value of regression parameter is only given for the intercept. The most parsimonious model is highlighted in bold font.

| Intercept | sex | stress | sex:stress | adj.$R^2$ | df | logLik | AICc | ΔAICc | weight |
| --- | --- | --- | --- | --- | --- | --- | --- | --- | --- |
| 7,91 | + | + | + | 0,25 | 5 | -29,17 | 69,50 | 0,00 | 0,32 |
| 7,99 | + | + |  | 0,20 | 4 | -30,45 | 69,66 | 0,17 | 0,30 |
| **7,91** |  | **+** |  | **0,15** | **3** | **-31,64** | **69,73** | **0,23** | **0,29** |
| 7,87 | + |  |  | 0,07 | 3 | -33,33 | 73,09 | 3,60 | 0,05 |
| 7,77 |  |  |  | 0,00 | 2 | -34,79 | 73,80 | 4,30 | 0,04 |

**Table S2j.** Effect of the water stress condition and sex on immune cell size. For each model, we reported intercept of the regression, adjusted R^2^ (adj.R^2^), degree of freedom (df), Log likelihood (LogLik) values, Akaike information criteria values with a correction for small sample sizes (AICc), change in AICc (ΔAICc) from the best model, and model weight. The presence of the categorial variable (sex, stress condition, and their interaction term sex:stress) in the model is indicated by a “+” symbol. The value of regression parameter is only given for the intercept. The most parsimonious model is highlighted in bold font.

| Intercept | sex | stress | sex:stress | adj.$R^2$ | df | logLik | AICc | ΔAICc | weight |
| --- | --- | --- | --- | --- | --- | --- | --- | --- | --- |
| 8,24 |  | + |  | 0,08 | 3 | -44,46 | 95,42 | 0,00 | 0,45 |
| **8,10** |  |  |  | **0,00** | **2** | **-46,36** | **96,97** | **1,54** | **0,21** |
| 8,17 | + | + |  | 0,10 | 4 | -44,10 | 97,06 | 1,63 | 0,20 |
| 8,04 | + |  |  | 0,01 | 3 | -46,16 | 98,81 | 3,39 | 0,08 |
| 8,19 | + | + | + | 0,10 | 5 | -44,06 | 99,42 | 4,00 | 0,06 |

#### Immune cell density

**Table S2k.** Effect of the thermal stress condition and sex on immune cell density. For each model, we reported intercept of the regression, adjusted R^2^ (adj.R^2^), degree of freedom (df), Log likelihood (LogLik) values, Akaike information criteria values with a correction for small sample sizes (AICc), change in AICc (ΔAICc) from the best model, and model weight. The presence of the categorial variable (sex, stress condition, and their interaction term sex:stress) in the model is indicated by a “+” symbol. The value of regression parameter is only given for the intercept. The most parsimonious model is highlighted in bold font.

| Intercept | sex | stress | sex:stress | adj.$R^2$ | df | logLik | AICc | ΔAICc | weight |
| --- | --- | --- | --- | --- | --- | --- | --- | --- | --- |
| **3538275,86** |  | + |  | **0,41** | **3** | **-898,33** | **1803,10** | **0,00** | **0,66** |
| 3644762,62 | + | + |  | 0,41 | 4 | -898,12 | 1804,99 | 1,89 | 0,26 |
| 3600666,67 | + | + | + | 0,41 | 5 | -898,08 | 1807,31 | 4,22 | 0,08 |
| 2472758,62 |  |  |  | 0,00 | 2 | -913,43 | 1831,07 | 27,98 | 0,00 |
| 2707777,78 | + |  |  | 0,02 | 3 | -912,92 | 1832,29 | 29,20 | 0,00 |

**Table S2l.** Effect of the water stress condition and sex on immune cell density. For each model, we reported intercept of the regression, adjusted R^2^ (adj.R^2^), degree of freedom (df), Log likelihood (LogLik) values, Akaike information criteria values with a correction for small sample sizes (AICc), change in AICc (ΔAICc) from the best model, and model weight. The presence of the categorial variable (sex, stress condition, and their interaction term sex:stress) in the model is indicated by a “+” symbol. The value of regression parameter is only given for the intercept. The most parsimonious model is highlighted in bold font.

| Intercept | sex | stress | sex:stress | adj.$R^2$ | df | logLik | AICc | ΔAICc | weight |
| --- | --- | --- | --- | --- | --- | --- | --- | --- | --- |
| **1465517,24** |  | + |  | **0,13** | **3** | **-802,79** | **1612,09** | **0,00** | **0,52** |
| 1827142,86 | + | + | + | 0,18 | 5 | -801,23 | 1613,77 | 1,68 | 0,23 |
| 1575106,08 | + | + |  | 0,14 | 4 | -802,60 | 1614,06 | 1,97 | 0,20 |
| 1892500,00 |  |  |  | 0,00 | 2 | -806,49 | 1617,23 | 5,14 | 0,04 |
| 1960434,78 | + |  |  | 0,00 | 3 | -806,44 | 1619,38 | 7,29 | 0,01 |

#### β-Galactosidase activity

**Table S2m.** Effect of the thermal stress condition and sex on β-Galactosidase activity. For each model, we reported intercept of the regression, adjusted R^2^ (adj.R^2^), degree of freedom (df), Log likelihood (LogLik) values, Akaike information criteria values with a correction for small sample sizes (AICc), change in AICc (ΔAICc) from the best model, and model weight. The presence of the categorial variable (sex, stress condition, and their interaction term sex:stress) in the model is indicated by a “+” symbol. The value of regression parameter is only given for the intercept. The most parsimonious model is highlighted in bold font.

| Intercept | sex | stress | sex:stress | adj.$R^2$ | df | logLik | AICc | ΔAICc | weight |
| --- | --- | --- | --- | --- | --- | --- | --- | --- | --- |
| **12399,83** |  | + |  | **0,17** | **3** | **-556,53** | **1119,51** | **0,00** | **0,60** |
| 12988,70 | + | + |  | 0,18 | 4 | -556,14 | 1121,06 | 1,55 | 0,28 |
| 12487,80 | + | + | + | 0,19 | 5 | -555,81 | 1122,82 | 3,30 | 0,11 |
| 14534,95 |  |  |  | 0,00 | 2 | -561,86 | 1127,94 | 8,42 | 0,01 |
| 15052,70 | + |  |  | 0,01 | 3 | -561,62 | 1129,71 | 10,20 | 0,00 |

**Table S2n.** Effect of the water stress condition and sex on β-Galactosidase activity. For each model, we reported intercept of the regression, adjusted R^2^ (adj.R^2^), degree of freedom (df), Log likelihood (LogLik) values, Akaike information criteria values with a correction for small sample sizes (AICc), change in AICc (ΔAICc) from the best model, and model weight. The presence of the categorial variable (sex, stress condition, and their interaction term sex:stress) in the model is indicated by a “+” symbol. The value of regression parameter is only given for the intercept. The most parsimonious model is highlighted in bold font.

| Intercept | sex | stress | sex:stress | adj.$R^2$ | df | logLik | AICc | ΔAICc | weight |
| --- | --- | --- | --- | --- | --- | --- | --- | --- | --- |
| **10694,60** |  | + |  | **0,17** | **3** | **-477,96** | **962,43** | **0,00** | **0,67** |
| 10867,90 | + | + |  | 0,18 | 4 | -477,84 | 964,53 | 2,10 | 0,23 |
| 10955,89 | + | + | + | 0,18 | 5 | -477,79 | 966,89 | 4,47 | 0,07 |
| 9725,79 |  |  |  | 0,00 | 2 | -482,92 | 970,08 | 7,65 | 0,01 |
| 10087,84 | + |  |  | 0,01 | 3 | -482,55 | 971,60 | 9,17 | 0,01 |

## Supplementary file 3: Graphical representations of results per sex

### 1. Life history traits

#### 1.A. Survival

**Figure 1.A.1:**
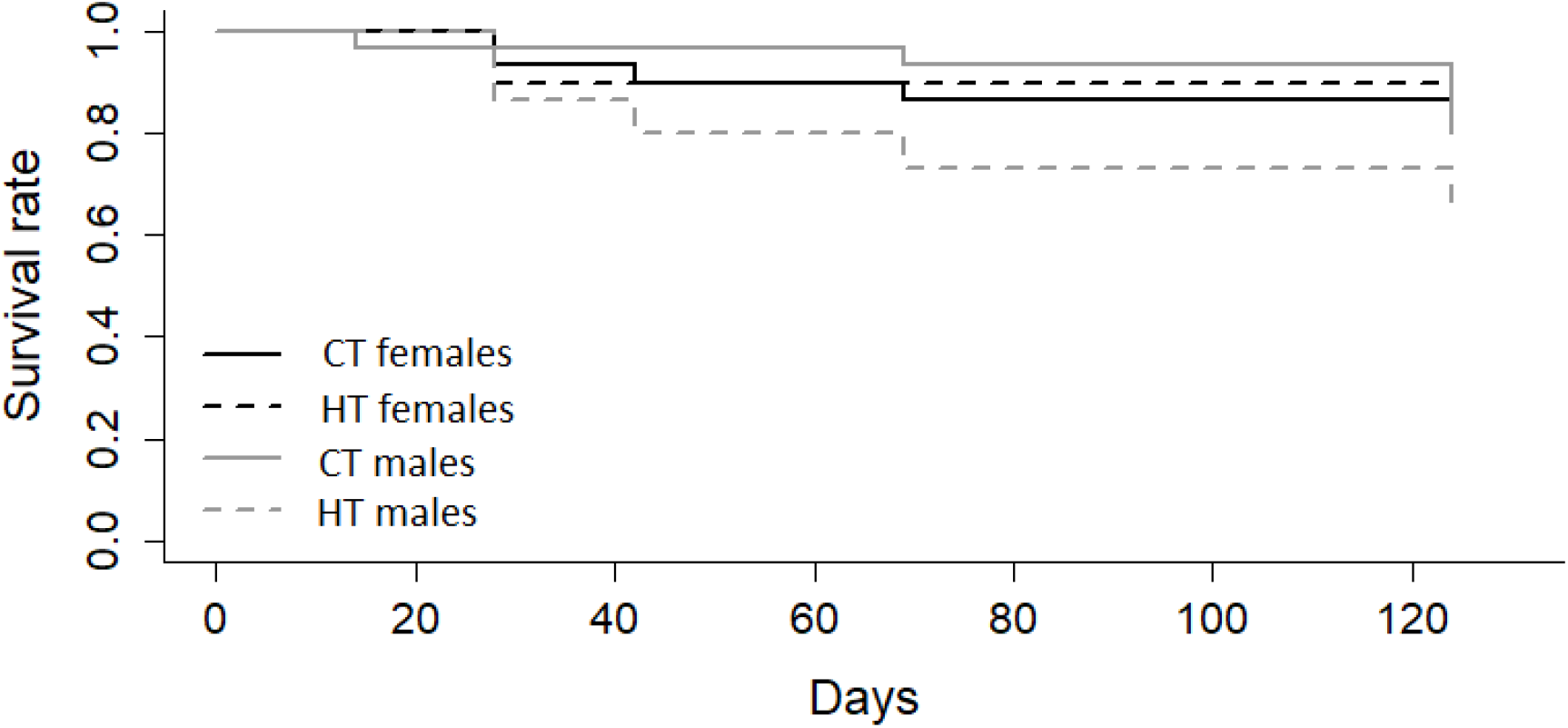
Effect of thermal stress on survival. CT females: control females in Control Temperature (20°C), HT females: stressed females in High Temperature (28°C), CT males: control males in Control Temperature (20°C), HT males: stressed males in High Temperature (28°C)

**Figure 1.A.2:**
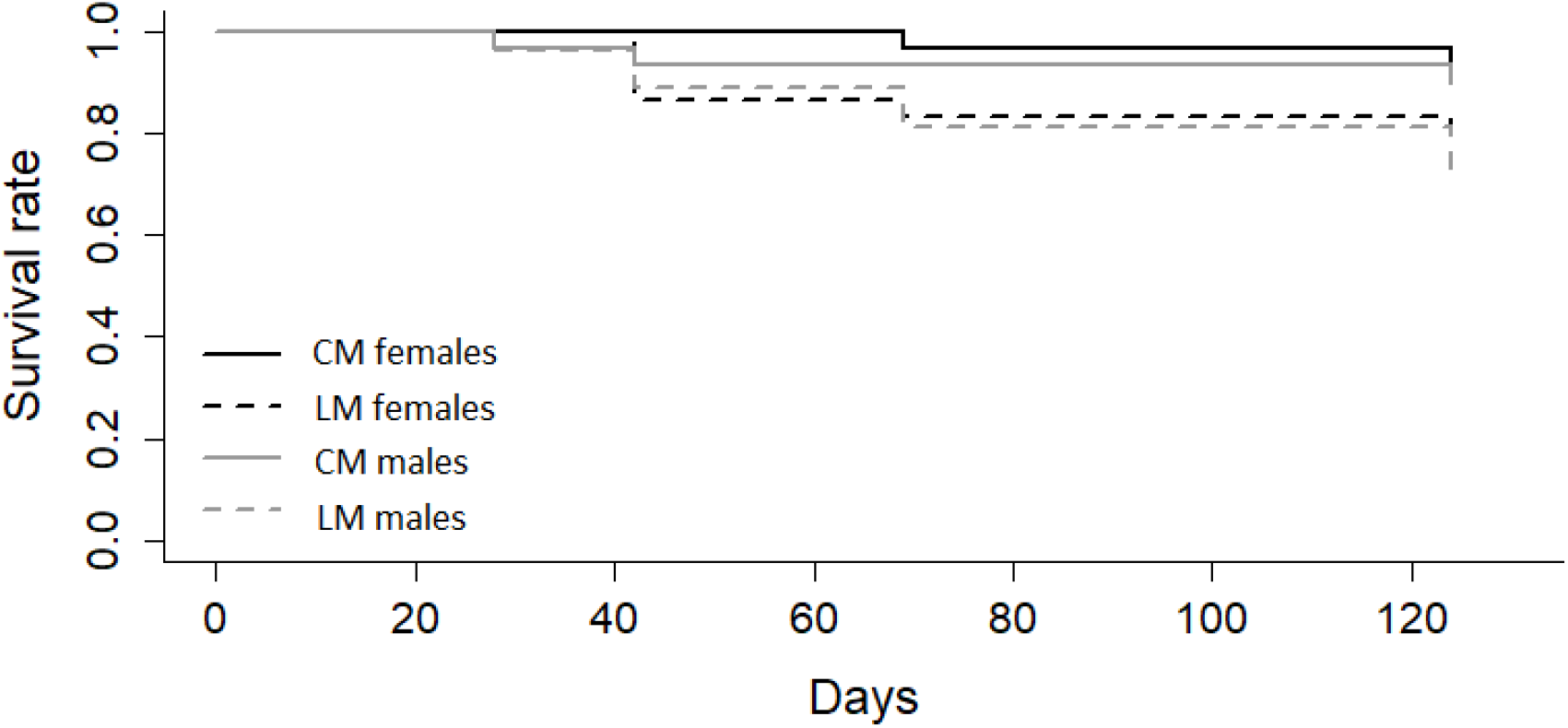
Effect of water stress on survival. CM females: control females in Control Moisture (moisture 80%), LM females: stressed females in Loss of Moisture (moisture 50%), CM males: control males in Control Moisture (moisture 80%), LM males: stressed males in Loss of Moisture (moisture 50%)

#### 1.B. Body mass across time

**Figure 1.B.1:**
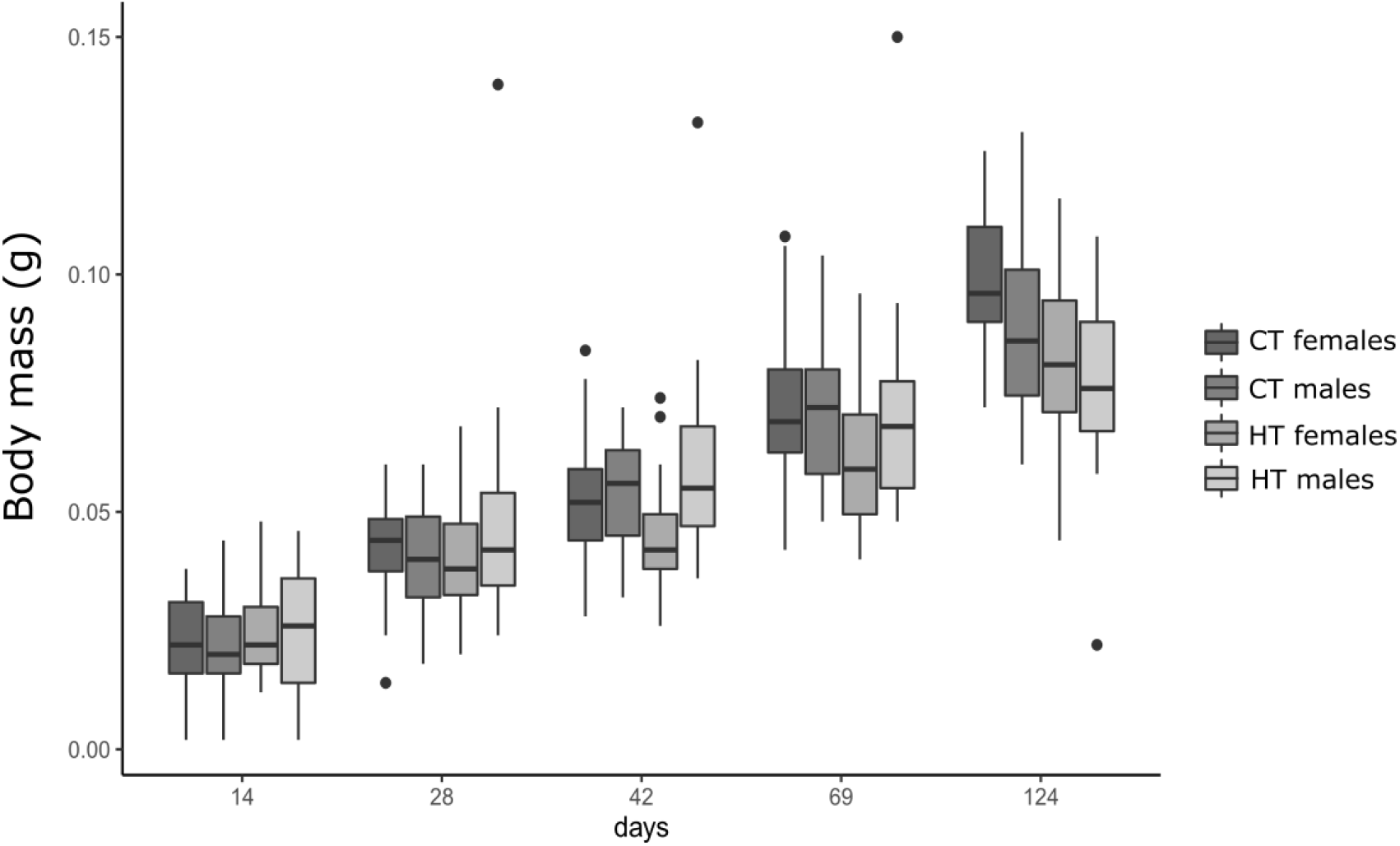
Boxplot of the effect of thermal stress on body mass (measured in grams) over time. CT females: control females in Control Temperature (20°C), HT females: stressed females in High Temperature (28°C), CT males: control males in Control Temperature (20°C), HT males: stressed males in High Temperature (28°C)

**Figure 1.B.2.:**
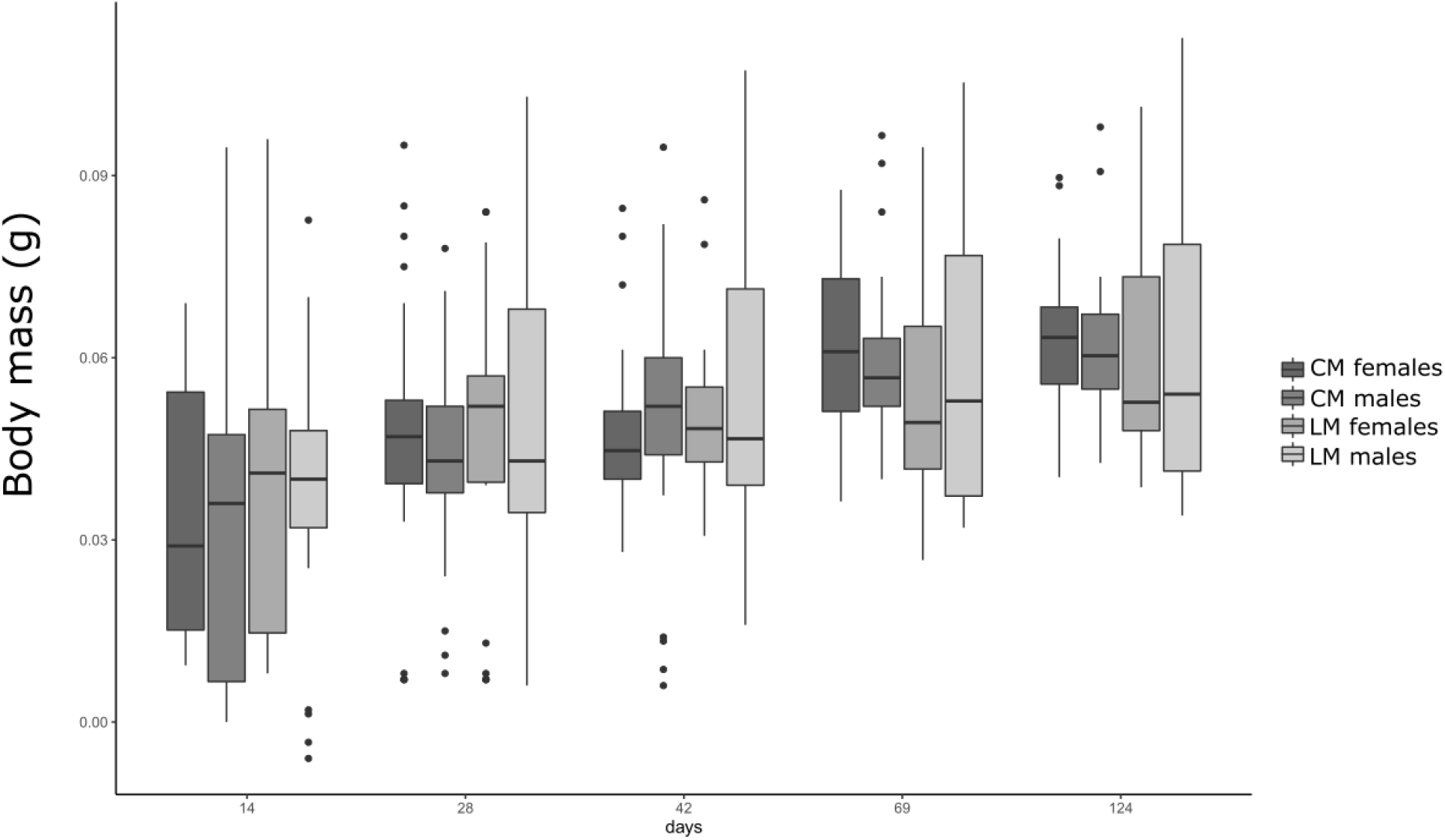
Boxplot of the effect of water stress on body mass (measured in grams) over time. CM females: control females in Control Moisture (moisture 80%), LM females: stressed females in Loss of Moisture (moisture 50%), CM males: control males in Control Moisture (moisture 80%), LM males: stressed males in Loss of Moisture (moisture 50%)

#### 1.C. Reproduction success

**Figure 1.C.1.:**
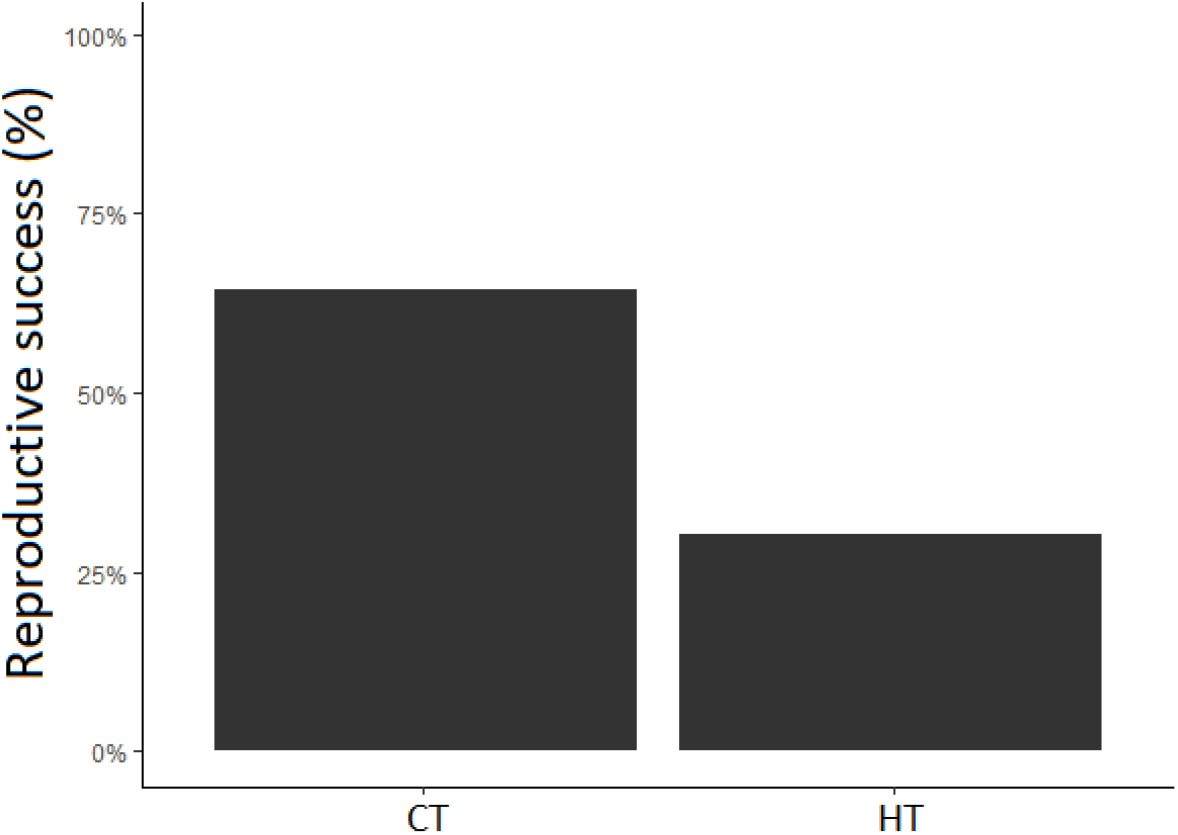
Effect of temperature on breeding success. (0 = pairs that did not produce offspring; 1 = pairs that produced offspring; CT: control individuals in Control Temperature (20°C), HT: Stressed individuals in High Temperature (28°C))

**Figure 1.C.2.:**
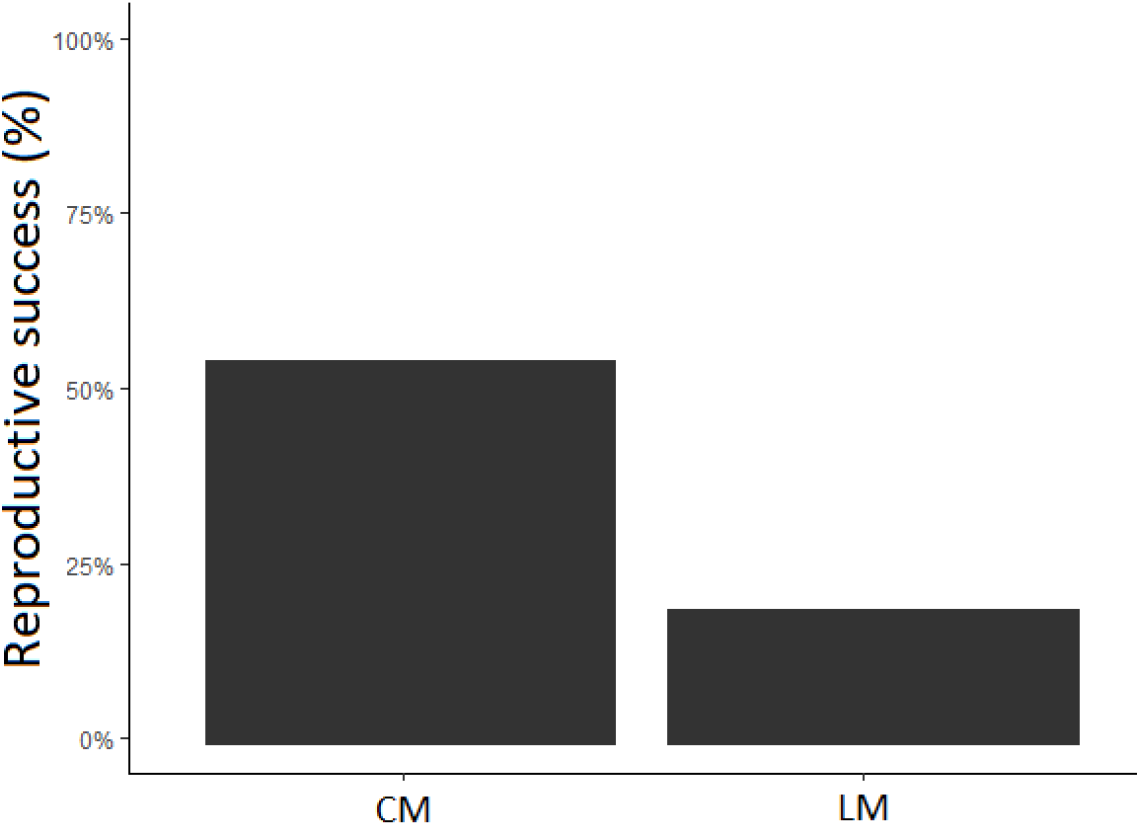
Effect of moisture on breeding success. (0 = pairs that did not produce offspring; 1 = pairs that produced offspring; CM: control individuals in Control Moisture (moisture 80%), LM females: stressed individuals in Loss of Moisture (moisture 50%))

### 2. Individual physiological traits

#### 2.A. Immune cells viability

**Figure 2.A.1.:**
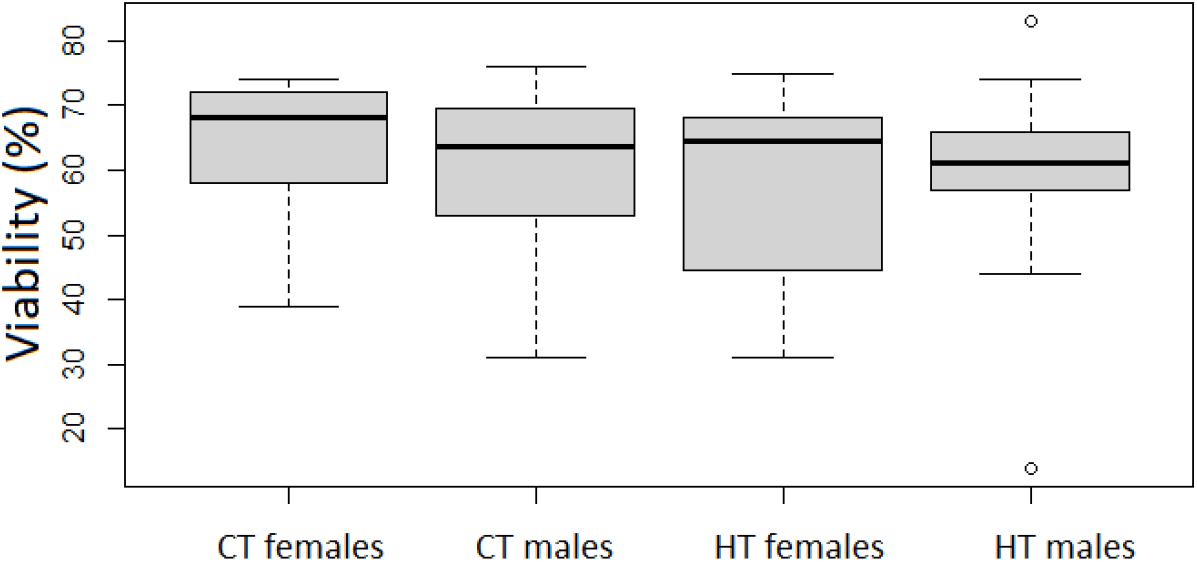
Effect of thermal stress on immune cell viability (% of live cells) CT females: control females in Control Temperature (20°C), HT females: stressed females in High Temperature (28°C), CT males: control males in Control Temperature (20°C), HT males: stressed males in High Temperature (28°C)

**Figure 2.A.2.:**
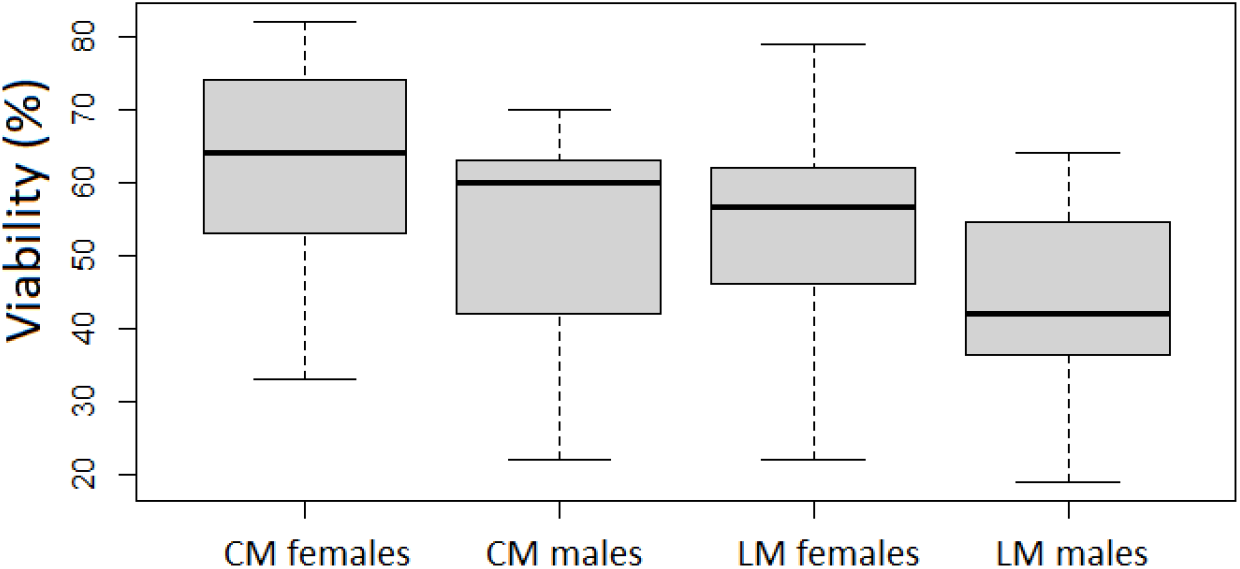
Effect of water stress on immune cell viability (% of live cells) CM females: control females in Control Moisture (moisture 80%), LM females: stressed females in Loss of Moisture (moisture 50%), CM males: control males in Control Moisture (moisture 80%), LM males: stressed males in Loss of Moisture (moisture 50%)

#### 2.B. Immune cells size

**Figure 2.B.1.:**
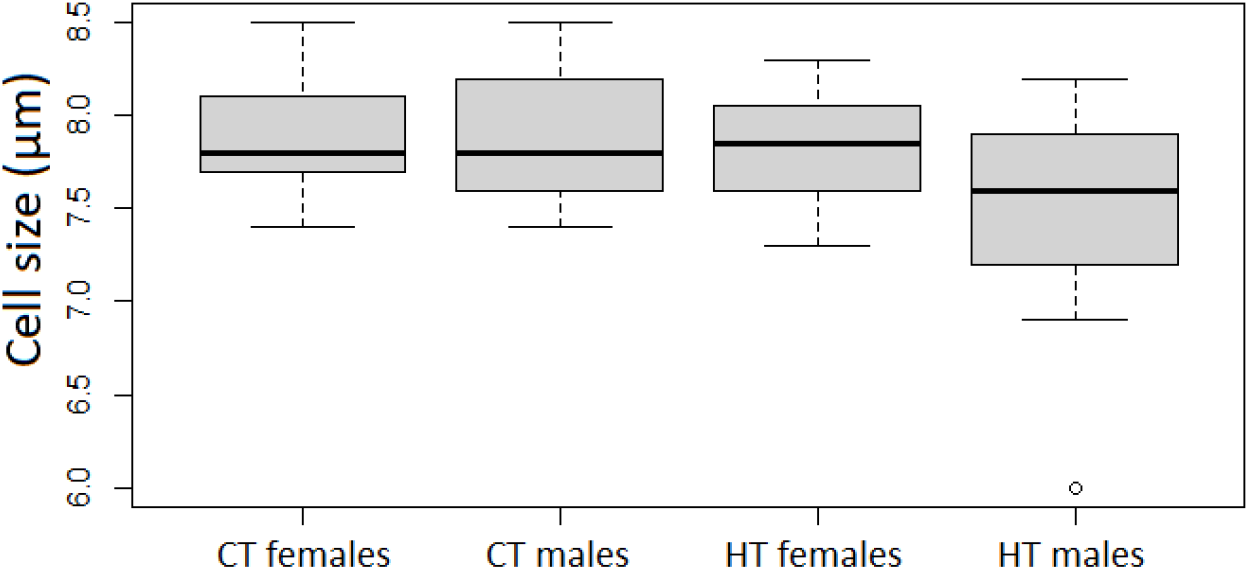
Effect of thermal stress on immune cells size (in μm) CT females: control females in Control Temperature (20°C), HT females: stressed females in High Temperature (28°C), CT males: control males in Control Temperature (20°C), HT males: stressed males in High Temperature (28°C)

**Figure 2.B.2.:**
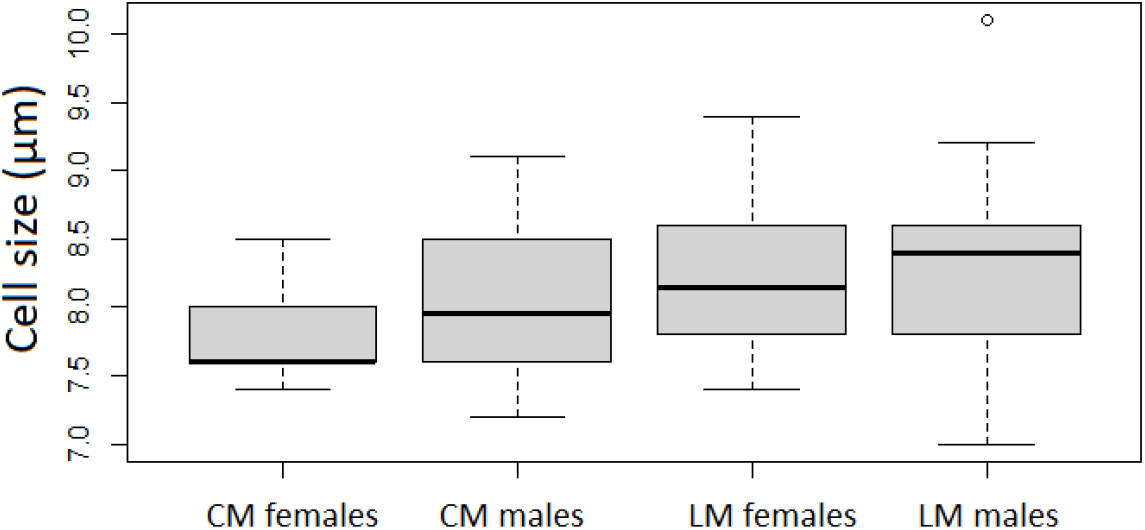
Effect of water stress on immune cells size (in μm) CM females: control females in Control Moisture (moisture 80%), LM females: stressed females in Loss of Moisture (moisture 50%), CM males: control males in Control Moisture (moisture 80%), LM males: stressed males in Loss of Moisture (moisture 50%)

#### 2.C. Immune cells density

**Figure 2.C.1.:**
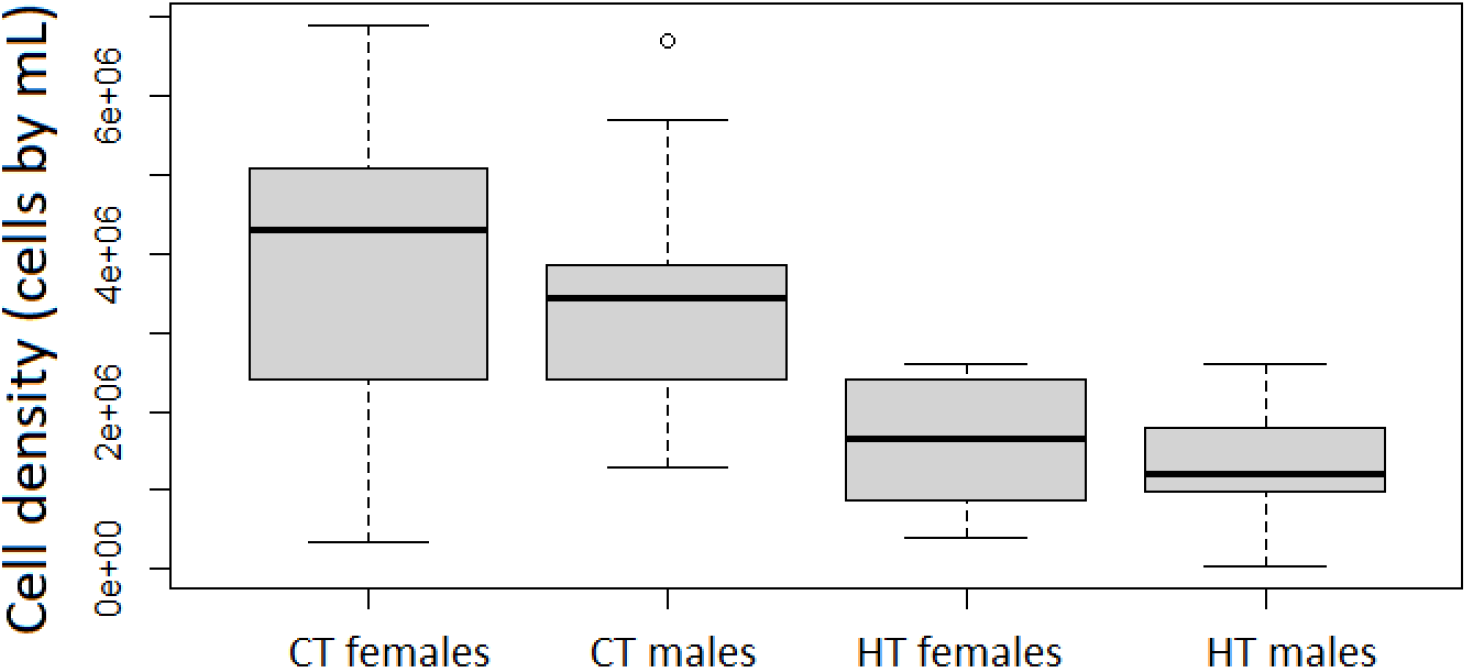
Effect of thermal stress on immune cells density (number of cells per mL of haemolymph) CT females: control females in Control Temperature (20°C), HT females: stressed females in High Temperature (28°C), CT males: control males in Control Temperature (20°C), HT males: stressed males in High Temperature (28°C)

**Figure 2.C.2.:**
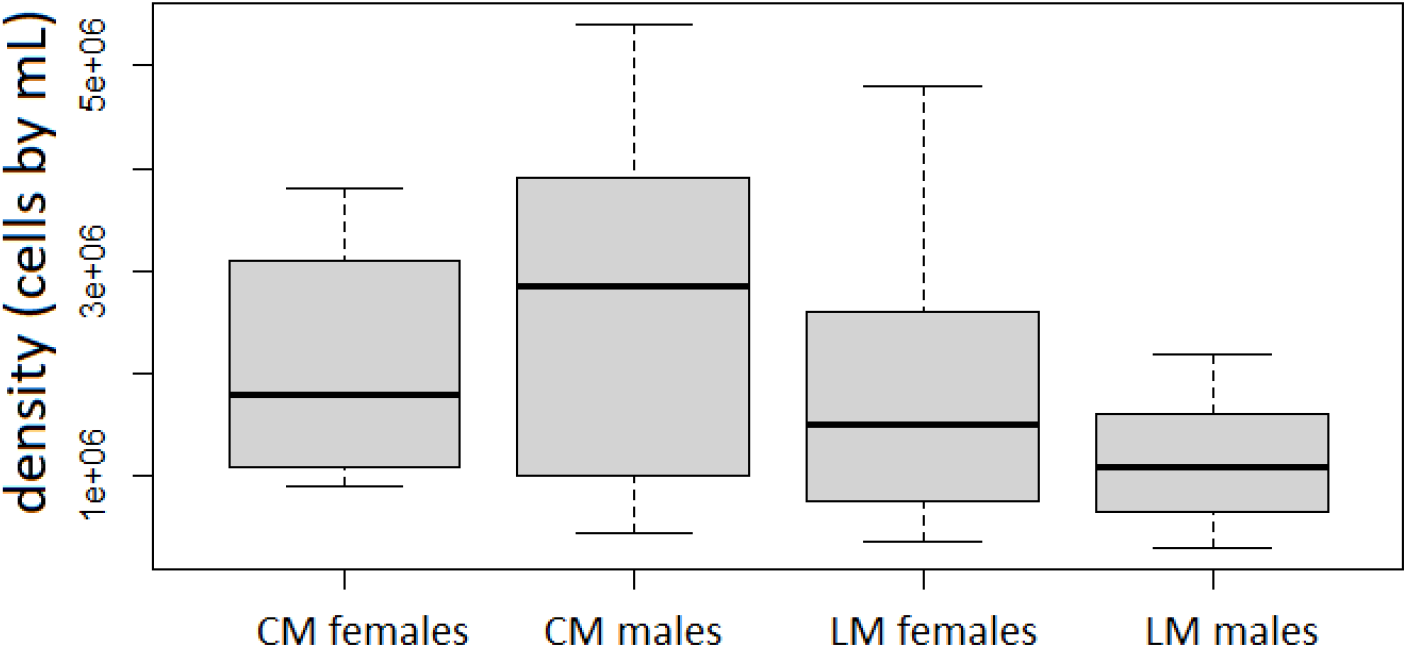
Effect of water stress on immune cells density (number of cells per mL of haemolymph) CM females: control females in Control Moisture (moisture 80%), LM females: stressed females in Loss of Moisture (moisture 50%), CM males: control males in Control Moisture (moisture 80%), LM males: stressed males in Loss of Moisture (moisture 50%)

#### 2.D. β-galactosidase activity

**Figure 2.D.1.:**
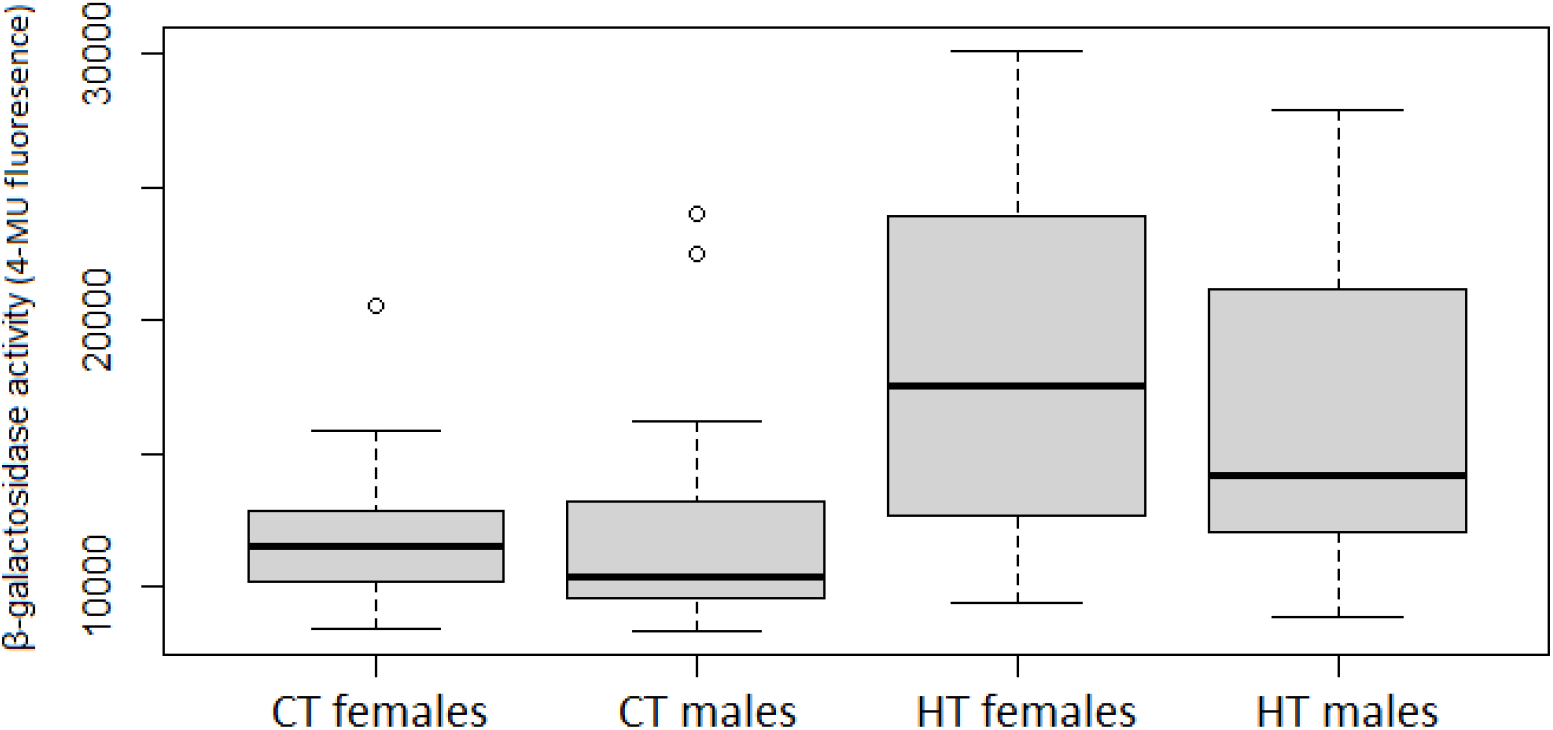
Effect of thermal stress on β-galactosidase activity. CT females: control females in Control Temperature (20°C), HT females: stressed females in High Temperature (28°C), CT males: control males in Control Temperature (20°C), HT males: stressed males in High Temperature (28°C)

**Figure 2.D.2.:**
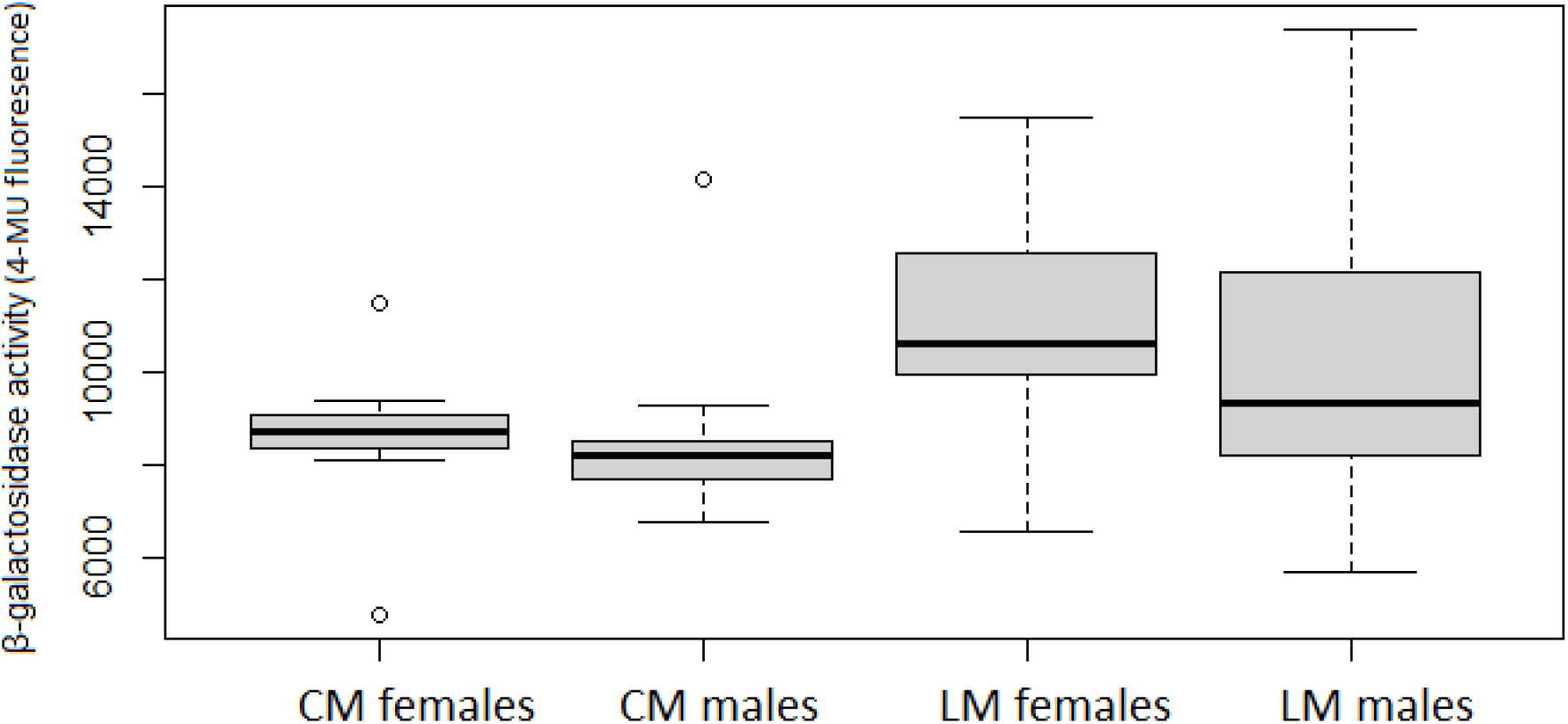
Effect of water stress on β-galactosidase activity. CM females: control females in Control Moisture (moisture 80%), LM females: stressed females in Loss of Moisture (moisture 50%), CM males: control males in Control Moisture (moisture 80%), LM males: stressed males in Loss of Moisture (moisture 50%)

