## Supplementary file 1 for "Deleterious effects of thermal and water stresses on life history and physiology: a case study on woodlouse"

**Supplementary File 1: Comparison of the two control groups**

Table 1: Comparison between the two control groups (CT (Control Temperature) and CM (Control Moisture)) of the two experiments for each tested variable (in bold the variables with significant statistical differences with graphical associated figures (Fig. 1, Fig. 2 and Fig. 3))

| **Traits** | **Statistical value** | **P-value** |
| --- | --- | --- |
| Life history traits measures |  |  |
| Survival | χ_1_² = 1.25 | P = 0.26 |
| **Body mass (day 14)** | **F_1,111_ = 13.00,** | **P < 0.001** |
| Reproduction | χ_1_² = 0.417 | P = 0.52 |
| Physiological traits measures |  |  |
| *Immune cells parameters* |  |  |
| **Density** | **F_1,50_ = 6.21** | **P= 0.016** |
| Viability | F_1,50_ = 2.49 | P= 0.12 |
| Size | F_1,50_ = 0.029 | P = 0.86 |
| **β-galactosidase activity** | F_1,49_ = 17.0 | **P < 0.001** |

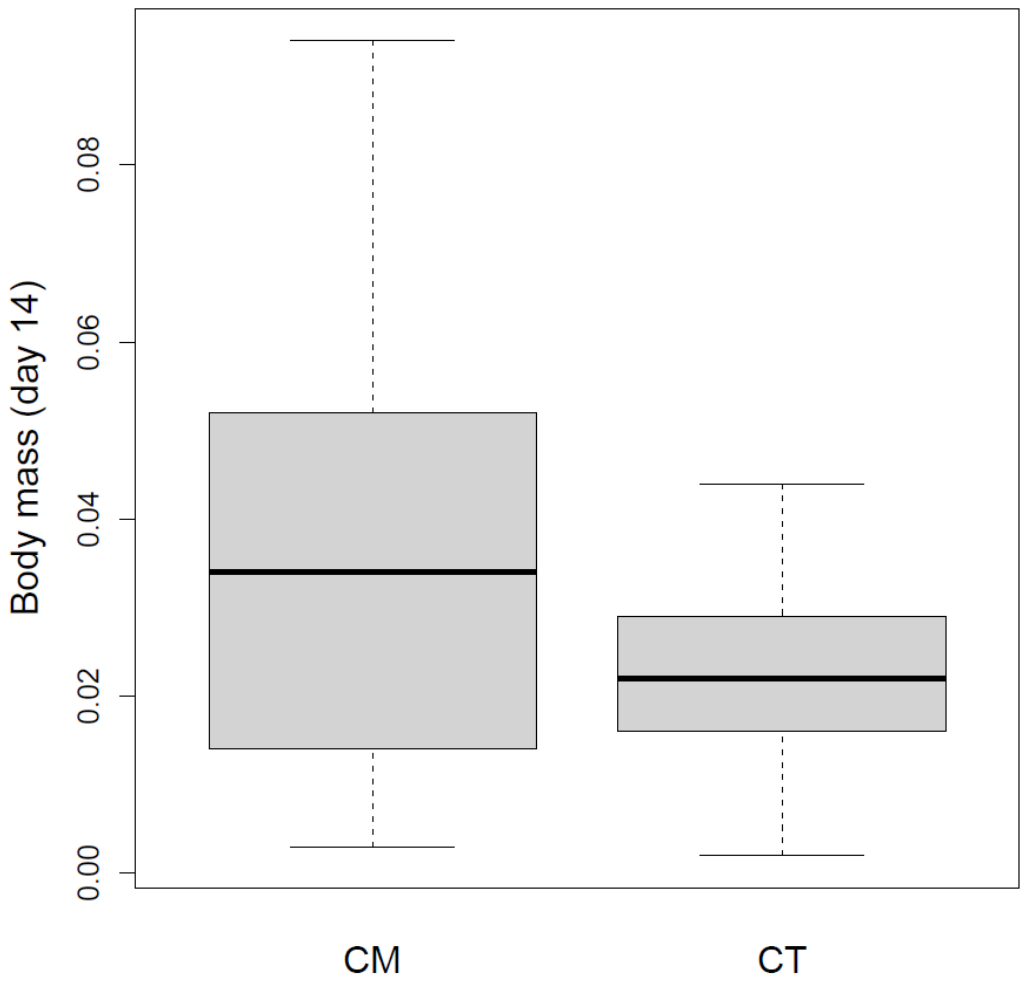

Figure 1: Body mass comparison between the two control groups

*(CT (Control Temperature) and CM (Control Moisture)) P-value < 0.001*

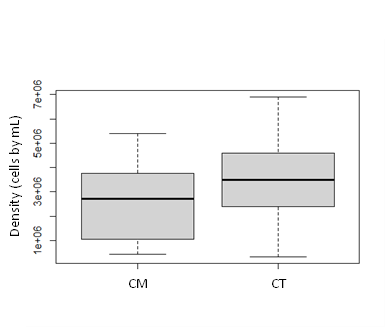

Figure 2: Immune cells density comparison between the two control groups

*(CT (Control Temperature) and CM (Control Moisture)) P-value=0.02*

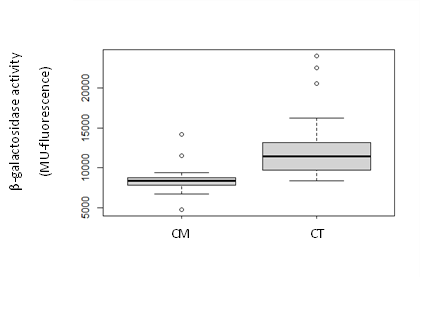

Figure 3: β-galactosidase activity comparison between the two control groups

*(CT (Control Temperature) and CM (Control Moisture))* *P-value<0.001*
