## Supplementary file 2 for "Deleterious effects of thermal and water stresses on life history and physiology: a case study on woodlouse"

**Supplementary File 2: Model selection**

**Life history trait**

**Survival**

**Table S2a.** Effect of the thermal stress condition and sex on the survival. For each model, we reported intercept of the regression, adjusted R² (adj.R²), degree of freedom (df), Log likelihood (LogLik) values, Akaike information criteria values with a correction for small sample sizes (AICc), change in AICc (△AICc) from the best model, and model weight. The presence of the categorial variable (sex, stress condition, and their interaction term sex:stress) in the model is indicated by a “+” symbol.

| Intercept | sex | stress | sex:stress | adj.R² | df | logLik | AICc | △AICc | weight |
| --- | --- | --- | --- | --- | --- | --- | --- | --- | --- |
| + |  | + |  | 0,08 | 1 | -125,01 | 252,18 | 0,00 | 0,31 |
| **+** |  |  |  | **0,00** | **0** | **-126,09** | **252,19** | **0,01** | **0,31** |
| + | + |  |  | 0,02 | 1 | -125,82 | 253,81 | 1,62 | 0,14 |
| + | + | + |  | 0,10 | 2 | -124,70 | 253,89 | 1,71 | 0,13 |
| + | + | + | + | 0,17 | 3 | -123,65 | 254,34 | 2,16 | 0,11 |

**Table S2b.** Effect of the water stress condition and sex on the survival. For each model, we reported intercept of the regression, adjusted R² (adj.R²), degree of freedom (df), Log likelihood (LogLik) values, Akaike information criteria values with a correction for small sample sizes (AICc), change in AICc (△AICc) from the best model, and model weight. The presence of the categorial variable (sex, stress condition, and their interaction term sex:stress) in the model is indicated by a “+” symbol. The most parsimonious model is highlighted in bold font.

| Intercept | sex | stress | sex:stress | adj.R² | df | logLik | AICc | △AICc | weight |
| --- | --- | --- | --- | --- | --- | --- | --- | --- | --- |
| **+** |  | **+** |  | **0,20** | **1** | **-91,25** | **184,72** | **0,00** | **0,53** |
| + | + | + |  | 0,22 | 2 | -91,02 | 186,75 | 2,02 | 0,19 |
| + |  |  |  | 0,00 | 0 | -93,52 | 187,04 | 2,32 | 0,17 |
| + | + |  |  | 0,02 | 1 | -93,32 | 188,87 | 4,15 | 0,07 |
| + | + | + | + | 0,23 | 3 | -90,94 | 189,39 | 4,67 | 0,05 |

**Body mass**

**Table S2c.** Effect of the thermal stress condition, sex and day on body mass. For each model, we reported intercept of the regression, adjusted R² (adj.R²), degree of freedom (df), Log likelihood (LogLik) values, Akaike information criteria values with a correction for small sample sizes (AICc), change in AICc (△AICc) from the best model, and model weight. The presence of the categorial variable (sex, stress condition, and their two-by-two interaction terms) in the model is indicated by a “+” symbol. The regression parameter is only given for the corresponding continuous variable (day) when this variable is present in the model. The most parsimonious model is highlighted in bold font.

| Intercept | day | sex | stress | day:sex | day:stress | sex:stress | adj.R² | df | logLik | AICc | △AICc | weight |
| --- | --- | --- | --- | --- | --- | --- | --- | --- | --- | --- | --- | --- |
| 0,02 | 0,00 | + | + | + | + | + | -0,01 | 8 | 1444,54 | -2872,80 | 0,00 | 0,63 |
| **0,02** | **0,00** | **+** | **+** |  | **+** | **+** | **-0,01** | **7** | **1442,55** | **-2870,90** | **1,91** | **0,24** |
| 0,02 | 0,00 | + | + | + | + |  | -0,01 | 7 | 1441,20 | -2868,18 | 4,62 | 0,06 |
| 0,02 | 0,00 |  | + |  | + |  | -0,01 | 5 | 1438,73 | -2867,35 | 5,45 | 0,04 |
| 0,02 | 0,00 | + | + |  | + |  | -0,01 | 6 | 1439,08 | -2866,01 | 6,79 | 0,02 |
| 0,02 | 0,00 | + | + | + |  | + | -0,01 | 7 | 1436,82 | -2859,43 | 13,37 | 0,00 |
| 0,03 | 0,00 | + | + |  |  | + | -0,01 | 6 | 1435,27 | -2858,39 | 14,41 | 0,00 |
| 0,02 | 0,00 | + | + | + |  |  | -0,01 | 6 | 1433,45 | -2854,75 | 18,05 | 0,00 |
| 0,03 | 0,00 |  | + |  |  |  | -0,01 | 4 | 1431,36 | -2854,64 | 18,16 | 0,00 |
| 0,02 | 0,00 | + | + |  |  |  | -0,01 | 5 | 1431,79 | -2853,47 | 19,33 | 0,00 |
| 0,02 | 0,00 | + |  | + |  |  | -0,01 | 5 | 1431,45 | -2852,78 | 20,02 | 0,00 |
| 0,02 | 0,00 |  |  |  |  |  | -0,01 | 3 | 1429,38 | -2852,71 | 20,09 | 0,00 |
| 0,02 | 0,00 | + |  |  |  |  | -0,01 | 4 | 1429,89 | -2851,71 | 21,09 | 0,00 |
| 0,05 |  |  | + |  |  |  | 0,00 | 3 | 1179,76 | -2353,47 | 519,33 | 0,00 |
| 0,05 |  |  |  |  |  |  | 0,00 | 2 | 1178,27 | -2352,52 | 520,28 | 0,00 |
| 0,05 |  | + | + |  |  |  | 0,00 | 4 | 1179,85 | -2351,62 | 521,18 | 0,00 |
| 0,06 |  | + | + |  |  | + | 0,00 | 5 | 1180,51 | -2350,91 | 521,90 | 0,00 |
| 0,05 |  | + |  |  |  |  | 0,00 | 3 | 1178,39 | -2350,74 | 522,06 | 0,00 |

**Table S2d.** Effect of the water stress condition, sex and day on body mass. For each model, we reported intercept of the regression, adjusted R² (adj.R²), degree of freedom (df), Log likelihood (LogLik) values, Akaike information criteria values with a correction for small sample sizes (AICc), change in AICc (△AICc) from the best model, and model weight. The presence of the categorial variable (sex, stress condition, and their two-by-two interaction terms) in the model is indicated by a “+” symbol. The regression parameter is only given for the corresponding continuous variable (day) when this variable is present in the model. The most parsimonious model is highlighted in bold font.

| Intercept | day | sex | stress | day:sex | day:stress | sex:stress | adj.R² | df | logLik | AICc | △AICc | weight |
| --- | --- | --- | --- | --- | --- | --- | --- | --- | --- | --- | --- | --- |
| **0,04** | **0,00** |  |  |  |  |  | **0,00** | **3** | **1309,82** | **-2613,60** | **0,00** | **0,29** |
| 0,04 | 0,00 |  | + |  | + |  | 0,00 | 5 | 1311,34 | -2612,57 | 1,03 | 0,17 |
| 0,04 | 0,00 | + |  |  |  |  | 0,00 | 4 | 1310,00 | -2611,93 | 1,68 | 0,12 |
| 0,04 | 0,00 |  | + |  |  |  | 0,00 | 4 | 1309,98 | -2611,89 | 1,72 | 0,12 |
| 0,04 | 0,00 | + | + |  | + |  | 0,00 | 6 | 1311,50 | -2610,85 | 2,76 | 0,07 |
| 0,04 | 0,00 | + | + |  |  |  | 0,00 | 5 | 1310,18 | -2610,24 | 3,37 | 0,05 |
| 0,04 | 0,00 | + |  | + |  |  | 0,00 | 5 | 1310,01 | -2609,90 | 3,70 | 0,04 |
| 0,04 | 0,00 | + | + |  | + | + | 0,00 | 7 | 1311,79 | -2609,37 | 4,24 | 0,03 |
| 0,04 | 0,00 | + | + | + | + |  | 0,00 | 7 | 1311,50 | -2608,79 | 4,81 | 0,03 |
| 0,04 | 0,00 | + | + |  |  | + | 0,00 | 6 | 1310,48 | -2608,79 | 4,82 | 0,03 |
| 0,04 | 0,00 | + | + | + |  |  | 0,00 | 6 | 1310,18 | -2608,21 | 5,40 | 0,02 |
| 0,04 | 0,00 | + | + | + | + | + | 0,00 | 8 | 1311,79 | -2607,30 | 6,30 | 0,01 |
| 0,04 | 0,00 | + | + | + |  | + | 0,00 | 7 | 1310,49 | -2606,75 | 6,85 | 0,01 |
| 0,05 |  |  |  |  |  |  | 0,00 | 2 | 1272,47 | -2540,92 | 72,68 | 0,00 |
| 0,05 |  | + |  |  |  |  | 0,00 | 3 | 1272,59 | -2539,13 | 74,47 | 0,00 |
| 0,05 |  |  | + |  |  |  | 0,00 | 3 | 1272,57 | -2539,10 | 74,50 | 0,00 |
| 0,05 |  | + | + |  |  |  | 0,00 | 4 | 1272,70 | -2537,32 | 76,28 | 0,00 |
| 0,05 |  | + | + |  |  | + | 0,00 | 5 | 1272,79 | -2535,47 | 78,13 | 0,00 |

**Reproductive success**

**Table S2e.** Effect of the thermal stress condition on the reproductive success. For each model, we reported intercept of the regression, adjusted R² (adj.R²), degree of freedom (df), Log likelihood (LogLik) values, Akaike information criteria values with a correction for small sample sizes (AICc), change in AICc (△AICc) from the best model, and model weight. The presence of the categorial variable (stress condition) in the model is indicated by a “+” symbol. The value of regression parameter is only given for the intercept. The most parsimonious model is highlighted in bold font.

| Intercept | stress | adj.R² | df | logLik | AICc | △AICc | weight |
| --- | --- | --- | --- | --- | --- | --- | --- |
| **0,62** | **+** | **0,16** | **2** | **-25,17** | **54,66** | **0,00** | **0,80** |
| -0,10 |  | 0,00 | 1 | -27,68 | 57,46 | 2,80 | 0,20 |

**Table S2f.** Effect of the water stress condition on the reproductive success. For each model, we reported intercept of the regression, adjusted R² (adj.R²), degree of freedom (df), Log likelihood (LogLik) values, Akaike information criteria values with a correction for small sample sizes (AICc), change in AICc (△AICc) from the best model, and model weight. The presence of the categorial variable (stress condition) in the model is indicated by a “+” symbol. The value of regression parameter is only given for the intercept. The most parsimonious model is highlighted in bold font.

| Intercept | stress | adj.R² | df | logLik | AICc | △AICc | weight |
| --- | --- | --- | --- | --- | --- | --- | --- |
| **-1,39** | **+** | **0,17** | **2** | **-23,77** | **51,87** | **0,00** | **0,83** |
| -0,51 |  | 0,00 | 1 | -26,46 | 55,03 | 3,16 | 0,17 |

**Individual physiological traits**

**Immune cell viability**

**Table** **S2g**. Effect of the thermal stress condition and sex on immune cell viability. For each model, we reported intercept of the regression, adjusted R² (adj.R²), degree of freedom (df), Log likelihood (LogLik) values, Akaike information criteria values with a correction for small sample sizes (AICc), change in AICc (△AICc) from the best model, and model weight. The presence of the categorial variable (sex, stress condition, and their interaction term sex:stress) in the model is indicated by a “+” symbol. The value of regression parameter is only given for the intercept. The most parsimonious model is highlighted in bold font.

| Intercept | sex | stress | sex:stress | adj.R² | df | logLik | AICc | △AICc | weight |
| --- | --- | --- | --- | --- | --- | --- | --- | --- | --- |
| **60,31** |  |  |  | **0,00** | **2** | **-231,04** | **466,30** | **0,00** | **0,47** |
| 61,97 |  | + |  | 0,02 | 3 | -230,57 | 467,58 | 1,28 | 0,25 |
| 61,00 | + |  |  | 0,00 | 3 | -230,97 | 468,39 | 2,08 | 0,17 |
| 62,43 | + | + |  | 0,02 | 4 | -230,53 | 469,81 | 3,51 | 0,08 |
| 63,67 | + | + | + | 0,03 | 5 | -230,24 | 471,63 | 5,32 | 0,03 |

**Table S2h.** Effect of the water stress condition and sex on immune cell viability. For each model, we reported intercept of the regression, adjusted R² (adj.R²), degree of freedom (df), Log likelihood (LogLik) values, Akaike information criteria values with a correction for small sample sizes (AICc), change in AICc (△AICc) from the best model, and model weight. The presence of the categorial variable (sex, stress condition, and their interaction term sex:stress) in the model is indicated by a “+” symbol. The value of regression parameter is only given for the intercept. The most parsimonious model is highlighted in bold font.

| Intercept | sex | stress | sex:stress | adj.R² | df | logLik | AICc | △AICc | weight |
| --- | --- | --- | --- | --- | --- | --- | --- | --- | --- |
| 51,82 | + | + |  | 0,15 | 4 | -212,23 | 433,32 | 0,00 | 0,48 |
| **47,45** |  | **+** |  | **0,08** | **3** | **-214,36** | **435,21** | **1,90** | **0,19** |
| 52,29 | + | + | + | 0,15 | 5 | -212,20 | 435,70 | 2,39 | 0,15 |
| 55,52 | + |  |  | 0,06 | 3 | -214,86 | 436,23 | 2,91 | 0,11 |
| 51,29 |  |  |  | 0,00 | 2 | -216,44 | 437,13 | 3,81 | 0,07 |

**Immune cell size**

**Table S2i.** Effect of the thermal stress condition and sex on immune cell size. For each model, we reported intercept of the regression, adjusted R² (adj.R²), degree of freedom (df), Log likelihood (LogLik) values, Akaike information criteria values with a correction for small sample sizes (AICc), change in AICc (△AICc) from the best model, and model weight. The presence of the categorial variable (sex, stress condition, and their interaction term sex:stress) in the model is indicated by a “+” symbol. The value of regression parameter is only given for the intercept. The most parsimonious model is highlighted in bold font.

| Intercept | sex | stress | sex:stress | adj.R² | df | logLik | AICc | △AICc | weight |
| --- | --- | --- | --- | --- | --- | --- | --- | --- | --- |
| 7,91 | + | + | + | 0,25 | 5 | -29,17 | 69,50 | 0,00 | 0,32 |
| 7,99 | + | + |  | 0,20 | 4 | -30,45 | 69,66 | 0,17 | 0,30 |
| **7,91** |  | **+** |  | **0,15** | **3** | **-31,64** | **69,73** | **0,23** | **0,29** |
| 7,87 | + |  |  | 0,07 | 3 | -33,33 | 73,09 | 3,60 | 0,05 |
| 7,77 |  |  |  | 0,00 | 2 | -34,79 | 73,80 | 4,30 | 0,04 |

**Table S2j.** Effect of the water stress condition and sex on immune cell size. For each model, we reported intercept of the regression, adjusted R² (adj.R²), degree of freedom (df), Log likelihood (LogLik) values, Akaike information criteria values with a correction for small sample sizes (AICc), change in AICc (△AICc) from the best model, and model weight. The presence of the categorial variable (sex, stress condition, and their interaction term sex:stress) in the model is indicated by a “+” symbol. The value of regression parameter is only given for the intercept. The most parsimonious model is highlighted in bold font.

| Intercept | sex | stress | sex:stress | adj.R² | df | logLik | AICc | △AICc | weight |
| --- | --- | --- | --- | --- | --- | --- | --- | --- | --- |
| 8,24 |  | + |  | 0,08 | 3 | -44,46 | 95,42 | 0,00 | 0,45 |
| **8,10** |  |  |  | **0,00** | **2** | **-46,36** | **96,97** | **1,54** | **0,21** |
| 8,17 | + | + |  | 0,10 | 4 | -44,10 | 97,06 | 1,63 | 0,20 |
| 8,04 | + |  |  | 0,01 | 3 | -46,16 | 98,81 | 3,39 | 0,08 |
| 8,19 | + | + | + | 0,10 | 5 | -44,06 | 99,42 | 4,00 | 0,06 |

**Immune cell density**

**Table S2k.** Effect of the thermal stress condition and sex on immune cell density. For each model, we reported intercept of the regression, adjusted R² (adj.R²), degree of freedom (df), Log likelihood (LogLik) values, Akaike information criteria values with a correction for small sample sizes (AICc), change in AICc (△AICc) from the best model, and model weight. The presence of the categorial variable (sex, stress condition, and their interaction term sex:stress) in the model is indicated by a “+” symbol. The value of regression parameter is only given for the intercept. The most parsimonious model is highlighted in bold font.

| Intercept | sex | stress | sex:stress | adj.R² | df | logLik | AICc | △AICc | weight |
| --- | --- | --- | --- | --- | --- | --- | --- | --- | --- |
| **3538275,86** |  | **+** |  | **0,41** | **3** | **-898,33** | **1803,10** | **0,00** | **0,66** |
| 3644762,62 | + | + |  | 0,41 | 4 | -898,12 | 1804,99 | 1,89 | 0,26 |
| 3600666,67 | + | + | + | 0,41 | 5 | -898,08 | 1807,31 | 4,22 | 0,08 |
| 2472758,62 |  |  |  | 0,00 | 2 | -913,43 | 1831,07 | 27,98 | 0,00 |
| 2707777,78 | + |  |  | 0,02 | 3 | -912,92 | 1832,29 | 29,20 | 0,00 |

**Table S2l.** Effect of the water stress condition and sex on immune cell density. For each model, we reported intercept of the regression, adjusted R² (adj.R²), degree of freedom (df), Log likelihood (LogLik) values, Akaike information criteria values with a correction for small sample sizes (AICc), change in AICc (△AICc) from the best model, and model weight. The presence of the categorial variable (sex, stress condition, and their interaction term sex:stress) in the model is indicated by a “+” symbol. The value of regression parameter is only given for the intercept. The most parsimonious model is highlighted in bold font.

| Intercept | sex | stress | sex:stress | adj.R² | df | logLik | AICc | △AICc | weight |
| --- | --- | --- | --- | --- | --- | --- | --- | --- | --- |
| **1465517,24** |  | **+** |  | **0,13** | **3** | **-802,79** | **1612,09** | **0,00** | **0,52** |
| 1827142,86 | + | + | + | 0,18 | 5 | -801,23 | 1613,77 | 1,68 | 0,23 |
| 1575106,08 | + | + |  | 0,14 | 4 | -802,60 | 1614,06 | 1,97 | 0,20 |
| 1892500,00 |  |  |  | 0,00 | 2 | -806,49 | 1617,23 | 5,14 | 0,04 |
| 1960434,78 | + |  |  | 0,00 | 3 | -806,44 | 1619,38 | 7,29 | 0,01 |

**β-Galactosidase activity**

**Table S2m.** Effect of the thermal stress condition and sex on β-Galactosidase activity. For each model, we reported intercept of the regression, adjusted R² (adj.R²), degree of freedom (df), Log likelihood (LogLik) values, Akaike information criteria values with a correction for small sample sizes (AICc), change in AICc (△AICc) from the best model, and model weight. The presence of the categorial variable (sex, stress condition, and their interaction term sex:stress) in the model is indicated by a “+” symbol. The value of regression parameter is only given for the intercept. The most parsimonious model is highlighted in bold font.

| Intercept | sex | stress | sex:stress | adj.R² | df | logLik | AICc | △AICc | weight |
| --- | --- | --- | --- | --- | --- | --- | --- | --- | --- |
| **12399,83** |  | **+** |  | **0,17** | **3** | **-556,53** | **1119,51** | **0,00** | **0,60** |
| 12988,70 | + | + |  | 0,18 | 4 | -556,14 | 1121,06 | 1,55 | 0,28 |
| 12487,80 | + | + | + | 0,19 | 5 | -555,81 | 1122,82 | 3,30 | 0,11 |
| 14534,95 |  |  |  | 0,00 | 2 | -561,86 | 1127,94 | 8,42 | 0,01 |
| 15052,70 | + |  |  | 0,01 | 3 | -561,62 | 1129,71 | 10,20 | 0,00 |

**Table S2n.** Effect of the water stress condition and sex on β-Galactosidase activity. For each model, we reported intercept of the regression, adjusted R² (adj.R²), degree of freedom (df), Log likelihood (LogLik) values, Akaike information criteria values with a correction for small sample sizes (AICc), change in AICc (△AICc) from the best model, and model weight. The presence of the categorial variable (sex, stress condition, and their interaction term sex:stress) in the model is indicated by a “+” symbol. The value of regression parameter is only given for the intercept. The most parsimonious model is highlighted in bold font.

| Intercept | sex | stress | sex:stress |  | adj.R² | df | logLik | AICc | △AICc | weight |
| --- | --- | --- | --- | --- | --- | --- | --- | --- | --- | --- |
| **10694,60** |  | **+** |  |  | **0,17** | **3** | **-477,96** | **962,43** | **0,00** | **0,67** |
| 10867,90 | + | + |  |  | 0,18 | 4 | -477,84 | 964,53 | 2,10 | 0,23 |
| 10955,89 | + | + | + |  | 0,18 | 5 | -477,79 | 966,89 | 4,47 | 0,07 |
| 9725,79 |  |  |  |  | 0,00 | 2 | -482,92 | 970,08 | 7,65 | 0,01 |
| 10087,84 | + |  |  |  | 0,01 | 3 | -482,55 | 971,60 | 9,17 | 0,01 |
