## Supplementary file 3 for "Deleterious effects of thermal and water stresses on life history and physiology: a case study on woodlouse"

**Supplementary file 3: Graphical representations of results per sex**

1. **Life history traits**

1.A. Survival
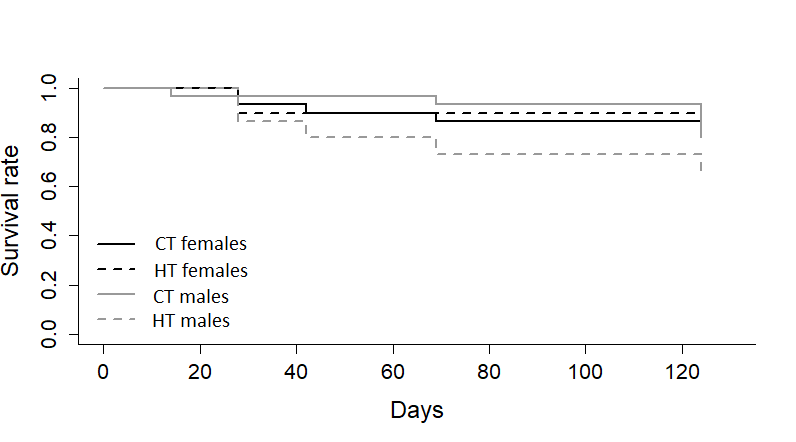

**Figure 1.A.1: Effect of thermal stress on survival**

*CT females: control females in Control Temperature (20°C), HT females: stressed females in High Temperature (28°C), CT males: control males in Control Temperature (20°C), HT males: stressed males in High Temperature (28°C)*

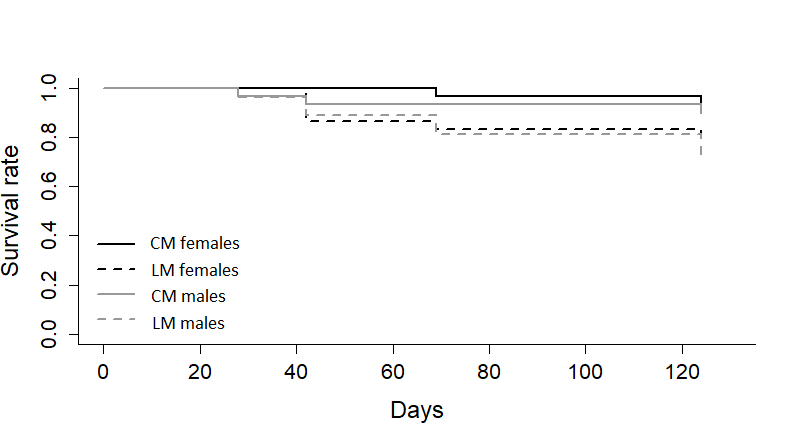

**Figure 1.A.2.: Effect of water stress on survival**

*CM females: control females in Control Moisture (moisture 80%), LM females: stressed females in Loss of Moisture (moisture 50%), CM males: control males in Control Moisture (moisture 80%), LM males: stressed males in Loss of Moisture (moisture 50%)*

1.B. Body mass across time

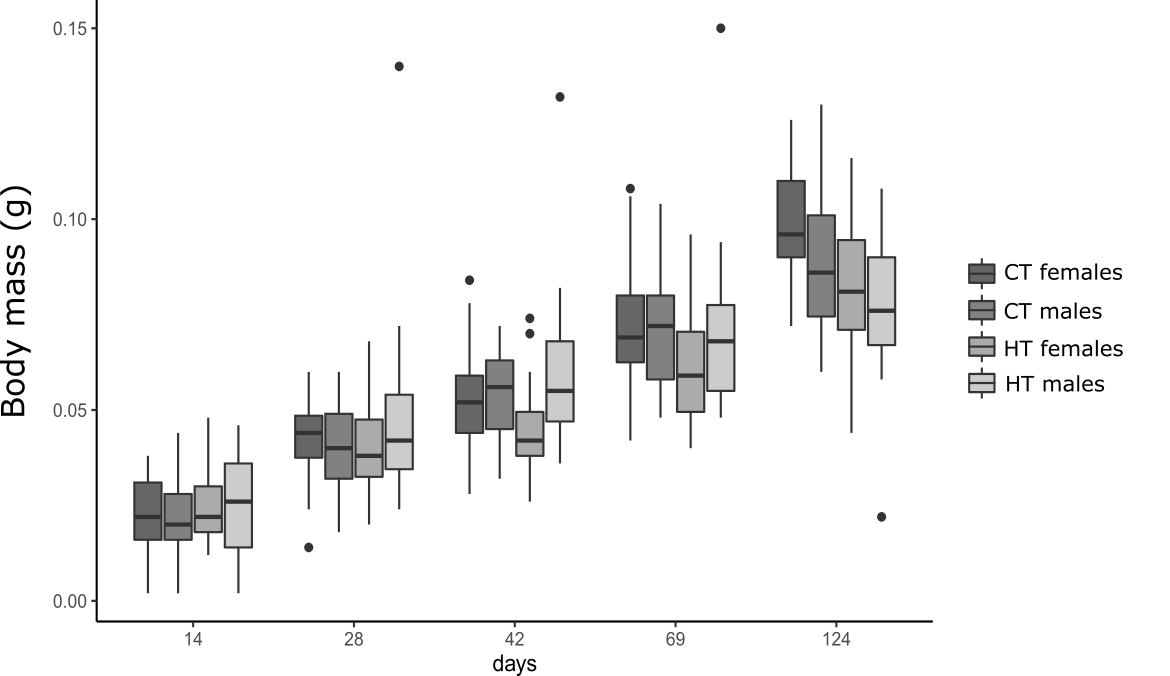

**Figure 1.B.1.: Boxplot of the effect of thermal stress on body mass (measured in grams) over time**

*CT females: control females in Control Temperature (20°C), HT females: stressed females in High Temperature (28°C), CT males: control males in Control Temperature (20°C), HT males: stressed males in High Temperature (28°C)*

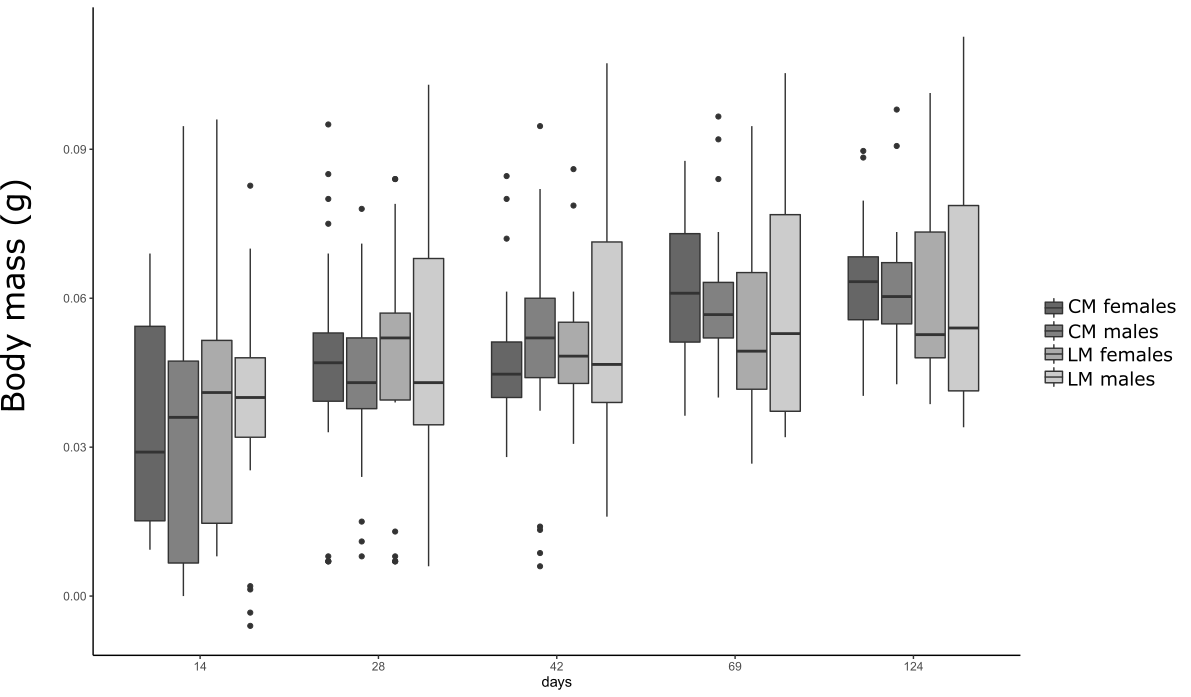

**Figure 1.B.2.: Boxplot of the effect of water stress on body mass (measured in grams) over time**

*CM females: control females in Control Moisture (moisture 80%), LM females: stressed females in Loss of Moisture (moisture 50%), CM males: control males in Control Moisture (moisture 80%), LM males: stressed males in Loss of Moisture (moisture 50%)*

1.C. Reproduction success

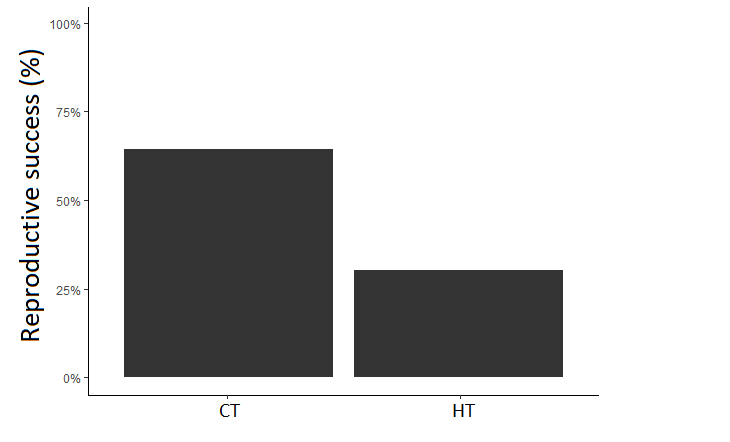

**Figure 1.C.1.: Effect of temperature on breeding success** (0 = pairs that did not produce offspring; 1 = pairs that produced offspring; CT: control individuals in Control Temperature (20°C), HT: Stressed individuals in High Temperature (28°C))

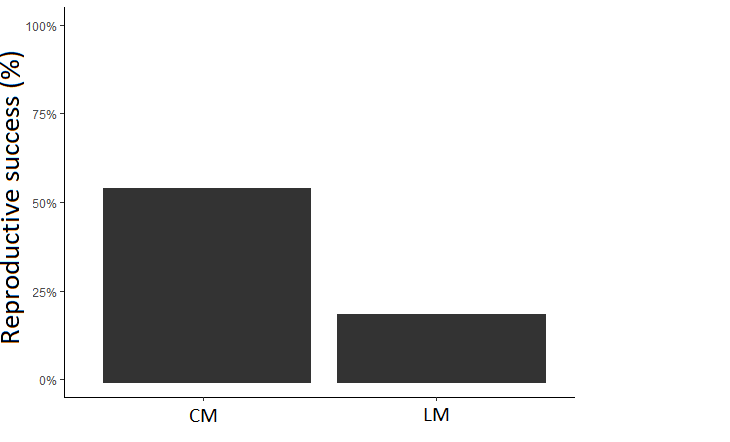

**Figure 1.C.2.: Effect of moisture on breeding success** (0 = pairs that did not produce offspring; 1 = pairs that produced offspring; CM: control individuals in Control Moisture (moisture 80%), LM females: stressed individuals in Loss of Moisture (moisture 50%))

2. **Individual physiological traits**

**2.A.** Immune cells viability

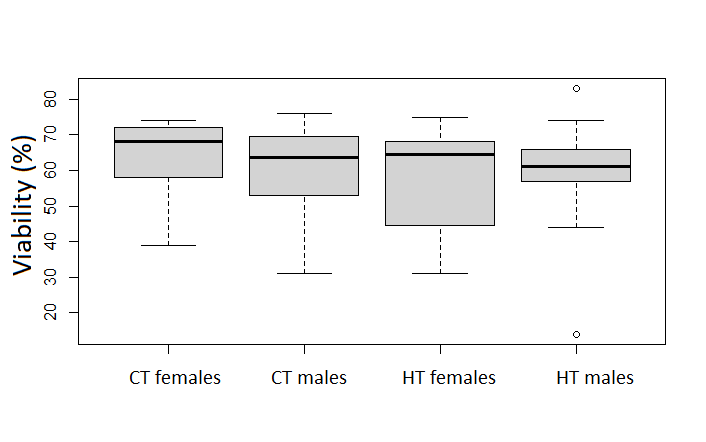

**Figure 2.A.1.: Effect of thermal stress on immune cell viability (% of live cells)**

*CT females: control females in Control Temperature (20°C), HT females: stressed females in High Temperature (28°C), CT males: control males in Control Temperature (20°C), HT males: stressed males in High Temperature (28°C)*

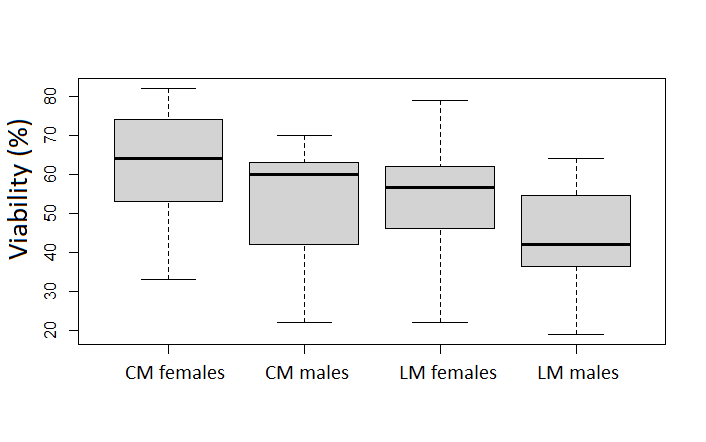

**Figure 2.A.2.: Effect of water stress on immune cell viability (% of live cells)**

*CM females: control females in Control Moisture (moisture 80%), LM females: stressed females in Loss of Moisture (moisture 50%), CM males: control males in Control Moisture (moisture 80%), LM males: stressed males in Loss of Moisture (moisture 50%)*

**2.B.** Immune cells size

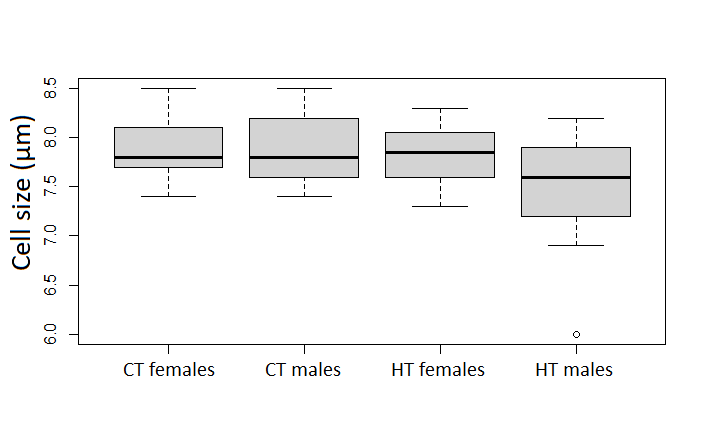

**Figure 2.B.1.: Effect of thermal stress on immune cells size (in µm)**

*CT females: control females in Control Temperature (20°C), HT females: stressed females in High Temperature (28°C), CT males: control males in Control Temperature (20°C), HT males: stressed males in High Temperature (28°C)*

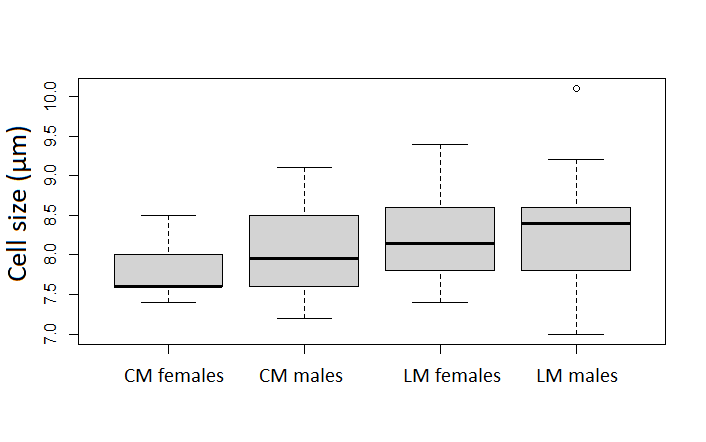

**Figure 2.B.2.: Effect of water stress on immune cells size (in µm)**

*CM females: control females in Control Moisture (moisture 80%), LM females: stressed females in Loss of Moisture (moisture 50%), CM males: control males in Control Moisture (moisture 80%), LM males: stressed males in Loss of Moisture (moisture 50%)*

**2.C.** Immune cells density

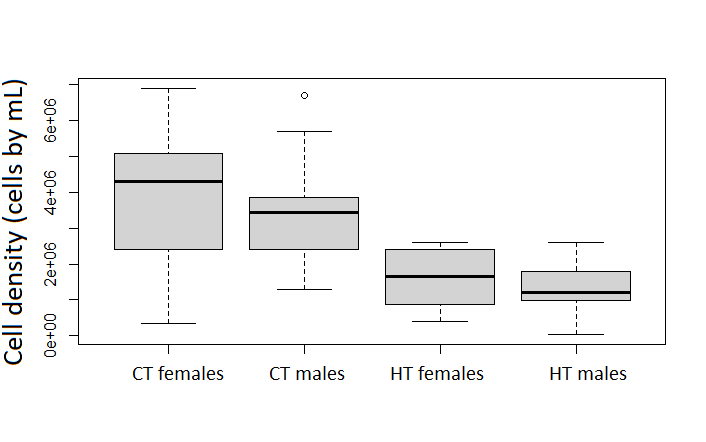

**Figure 2.C.1.: Effect of thermal stress on immune cells density (number of cells per mL of haemolymph)**

*CT females: control females in Control Temperature (20°C), HT females: stressed females in High Temperature (28°C), CT males: control males in Control Temperature (20°C), HT males: stressed males in High Temperature (28°C)*

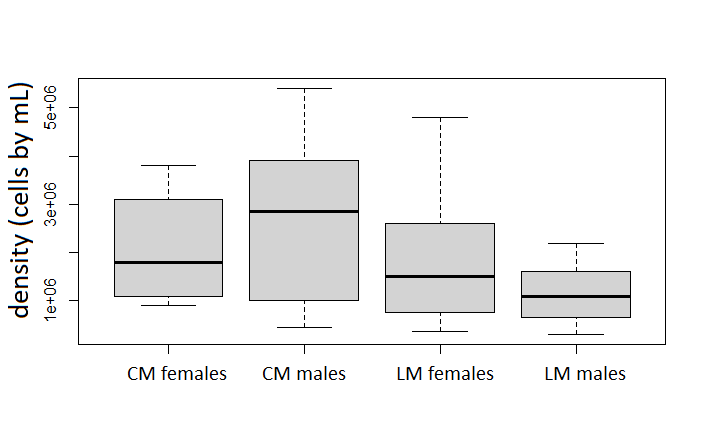

**Figure 2.C.2.: Effect of water stress on immune cells density (number of cells per mL of haemolymph)**

*CM females: control females in Control Moisture (moisture 80%), LM females: stressed females in Loss of Moisture (moisture 50%), CM males: control males in Control Moisture (moisture 80%), LM males: stressed males in Loss of Moisture (moisture 50%)*

**2.D**. β-galactosidase activity

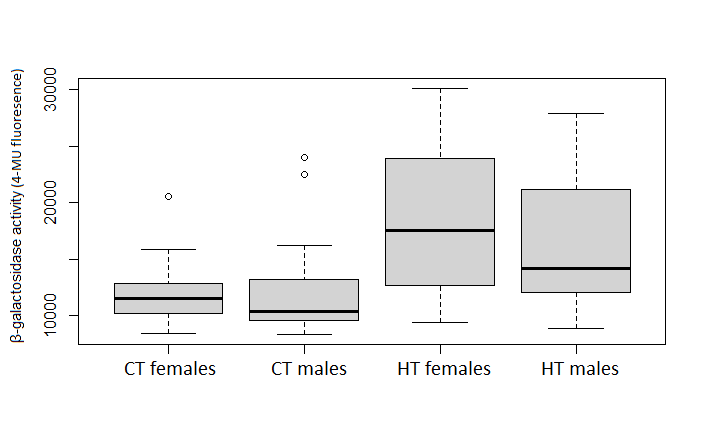

**Figure 2.D.1.: Effect of thermal stress on β-galactosidase activity**

*CT females: control females in Control Temperature (20°C), HT females: stressed females in High Temperature (28°C), CT males: control males in Control Temperature (20°C), HT males: stressed males in High Temperature (28°C)*

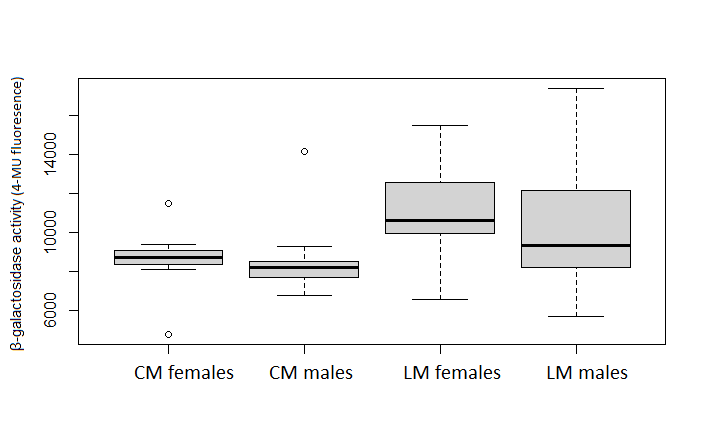

**Figure 2.D.2.: Effect of water stress on β-galactosidase activity**

*CM females: control females in Control Moisture (moisture 80%), LM females: stressed females in Loss of Moisture (moisture 50%), CM males: control males in Control Moisture (moisture 80%), LM males: stressed males in Loss of Moisture (moisture 50%)*
